# Structure of the human ATM kinase and mechanism of Nbs1 binding

**DOI:** 10.1101/2021.10.17.464701

**Authors:** C. Warren, N.P. Pavletich

## Abstract

DNA double-strand breaks (DSBs) can lead to mutations, chromosomal rearrangements, genome instability, and ultimately cancer. Central to the sensing of DSBs are ATM (Ataxia-telangiectasia mutated) kinase, which belongs to the phosphatidylinositol 3-kinase-related protein kinase (PIKK) family, and the MRN (Mre11-Rad50-Nbs1) protein complex that activates ATM. How the MRN complex recruits and activates ATM kinase is poorly understood. Previous studies indicate that the FxF/Y motif of Nbs1 directly binds to ATM kinase and is required to retain active ATM at sites of DNA damage. Here, we report the 2.5 Å resolution cryo-EM structures of human ATM and its complex with the Nbs1 FxF/Y motif. In keeping with previous structures of ATM and its yeast homolog Tel1, the dimeric human ATM kinase adopts a symmetric, butterfly-shaped autoinhibited structure. The conformation of the ATM kinase domain is most similar to the inactive states of other PIKKs, suggesting that activation may involve an analogous realigning the N and C lobes along with relieving the blockage of the substrate-binding site. We show that the Nbs1 FxF/Y motif binds to a conserved hydrophobic cleft within the Spiral domain of ATM, suggesting an allosteric mechanism of activation. We evaluate the importance of these interactions with mutagenesis and biochemical assays.

## Introduction

Ataxia-telangiectasia mutated (ATM) is a large protein kinase with a central role in the cellular response to DNA double-strand breaks (DSBs) and related genotoxic stress^1^. Mutations in ATM are responsible for Ataxia-telangiectasia (AT), which is a rare, autosomal recessive disorder characterized by telangiectasia, cerebellar degeneration, immunodeficiency, sensitivity to radiation and cancer susceptibility^2^.

ATM, which belongs to the phosphatidylinositol 3 kinase-like protein kinase (PIKK) family^2^, is essential for the sensing of DSBs during the cell cycle. It functions in association with the Mre11-Rad50-Nbs1 protein complex, termed MRN, which contributes to the localization of ATM to DSBs. The MRN complex, together with double-stranded DNA (dsDNA), serves to activate ATM as a protein kinase. ATM then phosphorylates a wide range of downstream effector proteins, such as p53, Chk2, Brca2, and H2A.X, leading to the activation of cell cycle checkpoints and homology-directed repair (HDR)^3, 4, 5–7^. Mutations in all three members of the MRN complex cause disorders that are phenotypically similar to AT^8–10^.

The MRN complex rapidly associates with DNA ends upon DSB formation, and it is a known component of ionizing radiation-induced foci (IRIF)^11, 12^. The Mre11 subunit has endonuclease and exonuclease activities that are implicated in DNA end processing^13, 14^, although nuclease-inactive mutants in yeast have only mild defects in responding to ionizing radiation (IR)^15^. Rad50 is a member of the diverse structural maintenance of chromatin (SMC) family of proteins. It contains N- and C-terminal ATPase domains separated by a long antiparallel coiled-coil with an apical conserved zinc hook that is thought to tether multiple MRN complexes^16–18^. Nbs1 contains N-terminal FHA and BRCT domains that are required for association with phosphorylated proteins such as H2A.X, CtIP, and Mdc1 at IRIF^19, 20^. While the C-terminal half of Nbs1 is largely disordered, it contains short sequence motifs for binding to Mre11 and ATM^21, 22^.

The activation of ATM by MRN is incompletely understood. The yeast homolog of ATM, Tel1, can associate with each of the individual subunits of the corresponding Mre11-Rad50-Xrs2 complex^23^. The Rad50 subunit plays an important role in activation, as either the Rad50-Mre11, or the Rad50-Xrs2 subcomplex can partially activate Tel1 in the presence of DNA, while the Mre11-Xrs2 complex cannot^23^. The ATM-binding segment of Nbs1 has been mapped to a sequence motif termed FxF/Y (residues 740-749) at the C-terminus^22^, however distinct ATM binding segments of Mre11 and Rad50 have not yet been identified. The Nbs1 FxF/Y motif is necessary for ATM recruitment to MRN-bound DSBs and for activation of the kinase in yeast and *Xenopus* extracts^22^. It is similarly required for ATM binding to MRN and for ATM recruitment to sites of DNA damage in human cell lines^24^. However, its deletion affects only a subset of the checkpoint functions, and it has non-uniform effects on downstream substrate phosphorylation^24^. In addition, fibroblasts from mice harboring an Nbs1 C-terminal deletion (Nbs1^ΔC/ΔC^) do not display overt sensitivity to IR, although they exhibit defective intra-S phase checkpoint activation, apoptosis induction, and phosphorylation of a subset of ATM targets^25^. Taken together, these results suggest that the ATM-Nbs1 interaction is not strictly required to initiate ATM activation, but it is likely necessary to stabilize activated ATM at sites of DNA damage and is critical for certain ATM-mediated checkpoint functions. This may reflect the ability of the remaining MRN subunits to also interact with ATM, or additional factors within IRIFs may aid in the recruitment of ATM *in vivo* to partially compensate for the loss of the Nbs1 C-terminus.

Along with ATM, the PIKK family also includes ATR, DNA-PKcs, mTOR, SMG1, and TRAPP^26^. These mammalian PIKKs coordinate diverse cellular processes, with ATR involved in the response to collapsed replication forks, DNA-PKcs in non-homologous end joining (NHEJ), mTOR in metabolism and cellular homeostasis, SMG1 in nonsense-mediated mRNA decay, and the catalytically inactive TRAPP in scaffolding chromatin remodeling complexes^27, 28^. Like ATM, these PIKK kinases are switched on by binding to their cognate activating proteins or protein complexes. The PIKKs share sequence homology across their C-terminal portion that consists of the ∼700-residue FAT domain (residues 1899-2613 of ATM) and subsequent ∼400-residue kinase domain (KD; residues 2614-3056 of ATM). Their N-terminal regions are divergent, and they adopt *α-a* solenoid structures typically consisting of two or more helical-repeat domains.

The cryo-EM structure of human ATM, based on a reconstruction of 4.4 Å to 5.7 Å resolution, showed that it forms a dimer through interactions between the FAT domains and also between the FAT and KD^29^. The FAT-KD interactions involve a region of the KD, termed the PRD (PIKK Regulatory Domain), that sequesters the putative polypeptide substrate-binding site^30^, as deduced from comparisons to canonical protein kinases^31^. This arrangement was thus suggested to maintain ATM in an inactive state by inhibiting substrate binding^29^. The N-terminal *α-a* solenoids are uninvolved in dimerization^29^. A subsequent cryo-EM analysis reported a monomeric form at 7.8 Å resolution, in addition to the canonical dimer at 4.3 Å^32^. The Tel1 homolog has been amenable to higher resolution structure determination^33^, with recent structures of the *Chaetomium thermophilum* and *Sacharomyces cerevisiae* Tel1 reported at overall resolutions of 3.7 Å and 3.9 Å, respectively^34, 35^. These Tel1 structures exhibited similar dimerization interfaces as ATM, including the region that blocks part of the putative substrate-binding site. Based on the higher resolutions of the Tel1 reconstructions, it was suggested that the kinase active site residues are in a catalytically competent organization, with the implication that the inaccessibility of the substrate-binding site is the primary mechanism of keeping ATM inactive. While Tel1 and ATM share a similar structural organization, the N-terminal ∼1800 residues preceding the FAT domain do not have detectable sequence homology.

Here we report the cryo-EM structure of human ATM at an overall resolution of 2.5 Å, as well as the structure of ATM bound to a peptide containing the Nbs1 FxF/Y motif. The refined model contains over 90 % of the ATM residues, with the remaining residues in poorly-ordered or disordered loops. The organization of residues in the ATM catalytic cleft are very similar to those of the inactive states of mTOR and DNA-PKcs, the two PIKKs for which high-resolution structures have been reported for both the inactive and active states^36–39^. Notably, ATM does not exhibit the conformational change that is characteristic of the active states of mTOR and DNA-PKcs kinase domains^37, 39^. Together with biochemical data, this suggests that activation of ATM by MRN may involve a conformational change that, in addition to relieving the partial blockage of the substrate-binding site, also realigns catalytic residues.

## Results

### Structure of the human ATM dimer

FLAG-tagged ATM was purified from a stably-transfected cell line (Supplemental Figure 1A). This preparation displays low but measurable kinase activity towards a p53 substrate peptide (Supplemental Figure 1B). Cryo-EM samples were prepared by mixing ATM with the non-hydrolyzable ATP analog adenylyl-imidodiphosphate (AMP-PNP) and MgCl_2_. The cryo-EM data yielded a consensus reconstruction in point group C2 that extended to an overall resolution of 2.5 Å as determined from gold standard Fourier shell correlation (FSC = 0.143) (Supplemental Table 1, Supplemental Figure 2). The maps show clear density for the majority of the side chains, allowing for the sequence assignment of the Pincer and Spiral domains and the accurate mapping of conserved residues and cancer-associated missense mutations (Supplemental Figures 3 to 7). While this manuscript was being prepared, a cryo-EM structure of ATM bound to the inhibitor KU-55933 was reported at an overall resolution of 2.8 Å^40^. Our structure of ATM bound to AMP-PNP is highly similar with a root mean square deviation (RMSD) of 0.9 Å based on 2453 aligned Cα atoms.

In keeping with previous structures of ATM and Tel1^29, 33–35, 41^, ATM adopts a butterfly shaped dimer with the FAT and KD domains (hereafter FATKD) forming a dimeric body and the N-terminal ∼1900 residues, previously described as Spiral and Pincer^29, 41^, extending away from this body. In our structure (Figure 1A to C), the N-terminal Spiral domain extends across residues 1 to 1166, the Pincer domain across residues 1167 to 1898, the FAT domain (named after FRAP, ATM, TRRAP) across residues 1899 to 2613, and the Kinase domain across residues 2614 to 3056. As described for other PIKKs^36, 37^, the Kinase domain consists of an N-terminal lobe (N lobe, residues 2614-2770) and a C-terminal lobe (C lobe, residues 2771-2957), with the catalytic cleft in between the two. The C lobe ends with the FAT C-terminal domain (FATC, residues 3027-3056) that is characteristic of the PIKK family and is absent from canonical kinases^36, 37^.

**Figure 1.**
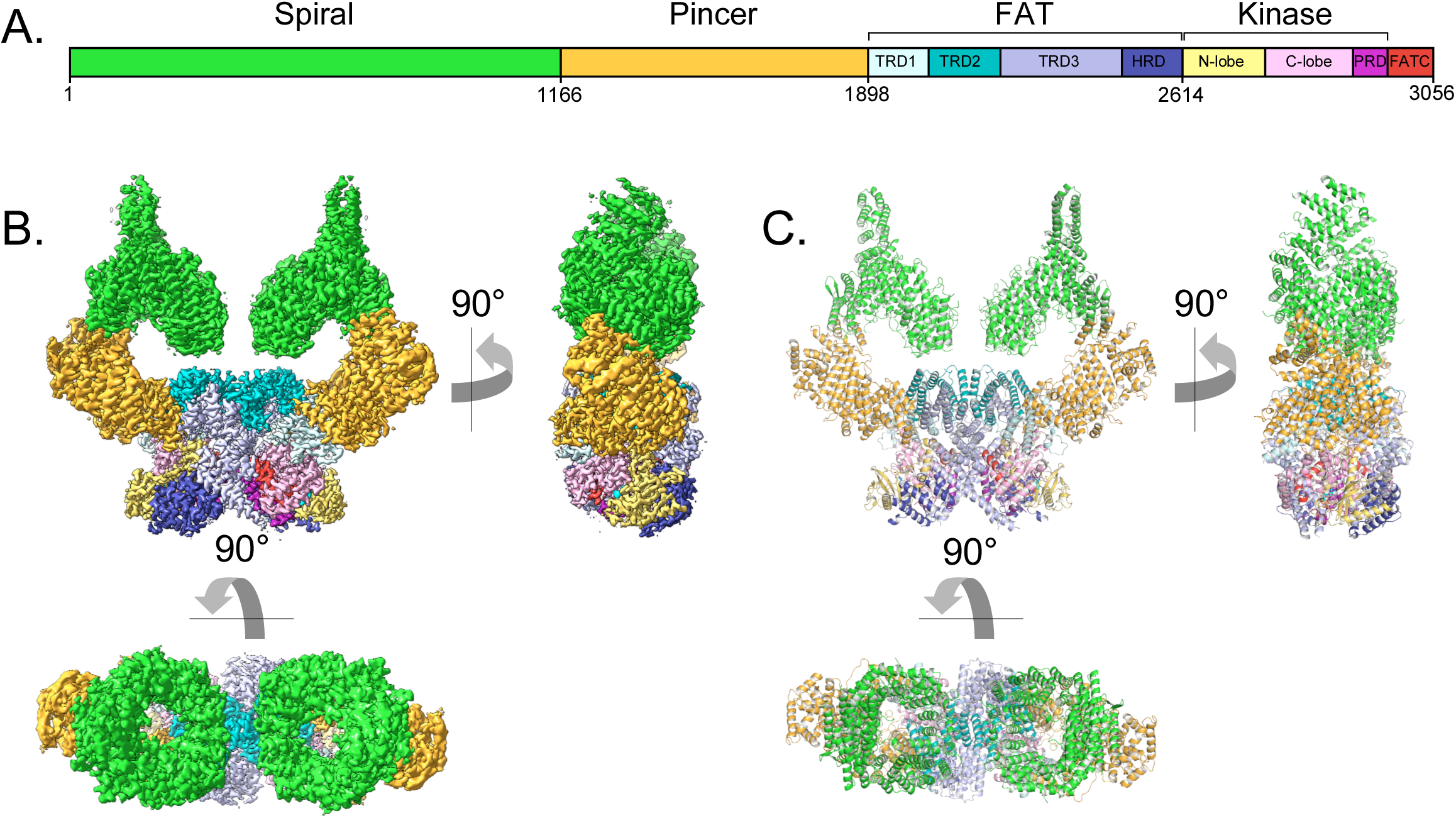
Overall structure of the human ATM kinase. **A)** Domain map of the human ATM kinase showing Spiral (residues 1-1166), Pincer (1167-1898), FAT (1899-2613), Kinase (2614-3026), and FATC (3027-3056) domains. FAT domain is further divided into TRD1 (1899-2025), TRD2 (2026-2192), TRD3 (2193-2479), and HRD (2480-2613) subdomains. Kinase domain is further divided into N-lobe (2614-2770), C-lobe (2771-2957), and PRD (2958-3026) subdomains. **B)** Composite cryo-EM density map of ATM kinase colored by approximate domain location. AMP-PNP molecule and Mg^2+^ ion in the active site colored cyan and gray, respectively. **C)** Structure of the overall ATM dimer colored by domain.

The N-terminal Spiral and Pincer domains, which consists mostly of HEAT repeats, are mobile relative to the FATKD dimer body. To improve the resolution of the reconstruction in this region, we used partial signal subtraction followed by symmetry expansion and focused refinement (see Methods and Supplemental Figure 3)^42^. The mobility originates in part from flexibility within the Pincer domain, which forms the elbow-like structure of the α-solenoid arm that extends from the N-terminus to the start of the FAT domain. This flexibility is associated with the solenoid arm adopting a continuum of positions relative to the FATKD. Similar, albeit more extensive flexibility has also been observed with the cryo-EM reconstruction of *C. thermophilum* Tel1^34^. Extensive 3D classification indicated that the Spiral and Pincer domains have no preferred conformation relative to the FATKD body (Supplemental Figure 2C). The most “open” and “closed” subclasses were each refined to an overall resolution of 2.8 Å as determined from the gold standard FSC (Supplemental Figure 2C and D). In the most open conformation, the N-terminal tips of the Spiral domains are separated by ∼134 Å, while these same regions are separated by ∼125 Å in the most closed conformation (Supplemental Figure 6A and B). Along with translational movement of these domains, we also observe a slight clockwise rotational motion upon closure, which is apparent when viewed along the symmetry axis of the enzyme (Supplemental Figure 6C and D, Supplemental Movie 1). The flexibility of the Spiral and Pincer domains does not lead to any significant conformational changes in the FATKD segment, which can be superimposed with a 0.39 Å RMSD in the positions of 1048 C*α* atoms from the open and closed conformation structures (Supplemental Figure 6E and F). Therefore, it is unlikely that any of these subclasses represent an intrinsically more active conformation of the ATM dimer.

The ATM FAT domain plays a central role in dimerization, which buries ∼3,800 Å^2^ of surface area on each ATM protomer (Supplemental Figure 7)^43^. As with other PIKKs, the FAT domain consists of three Tetratricopeptide Repeat Domains subdomains (TRD1 residues 1899-2025, TRD2 residues 2026-2192, and TRD3 residues 2193-2479) followed by a HEAT-repeat subdomain (HRD residues 2480-2613) (Figure 2A). It adopts a “C”- shaped structure that partially encircles the KD. In an arrangement that has been described as a C-clamp^36^, TRD1 packs with the KD C lobe, while the HRD packs with both N and C lobes adjacent to the catalytic cleft. This arrangement is conserved among structurally characterized PIKKs^26, 28^. TRD1 additionally packs with multiple regions of the Pincer domain that precedes it. Ser1981, a conserved TRD1 residue whose autophosphorylation coincides with ATM activation^44, 45^, is in a 9-residue disordered loop between the fα1c and fα1d *α* helices (prefix “f” denotes FAT domain helices) and is not visible in our map. However, due to the limited length of this loop and its position relative to the active site, it is unlikely that Ser1981 would be phosphorylated by either protomer in the dimeric structure. This suggests that Ser1981 is either phosphorylated by a separate ATM molecule during activation, or that a major MRN-induced structural rearrangement precedes Ser1981 autophosphorylation in the activation pathway.

**Figure 2.**
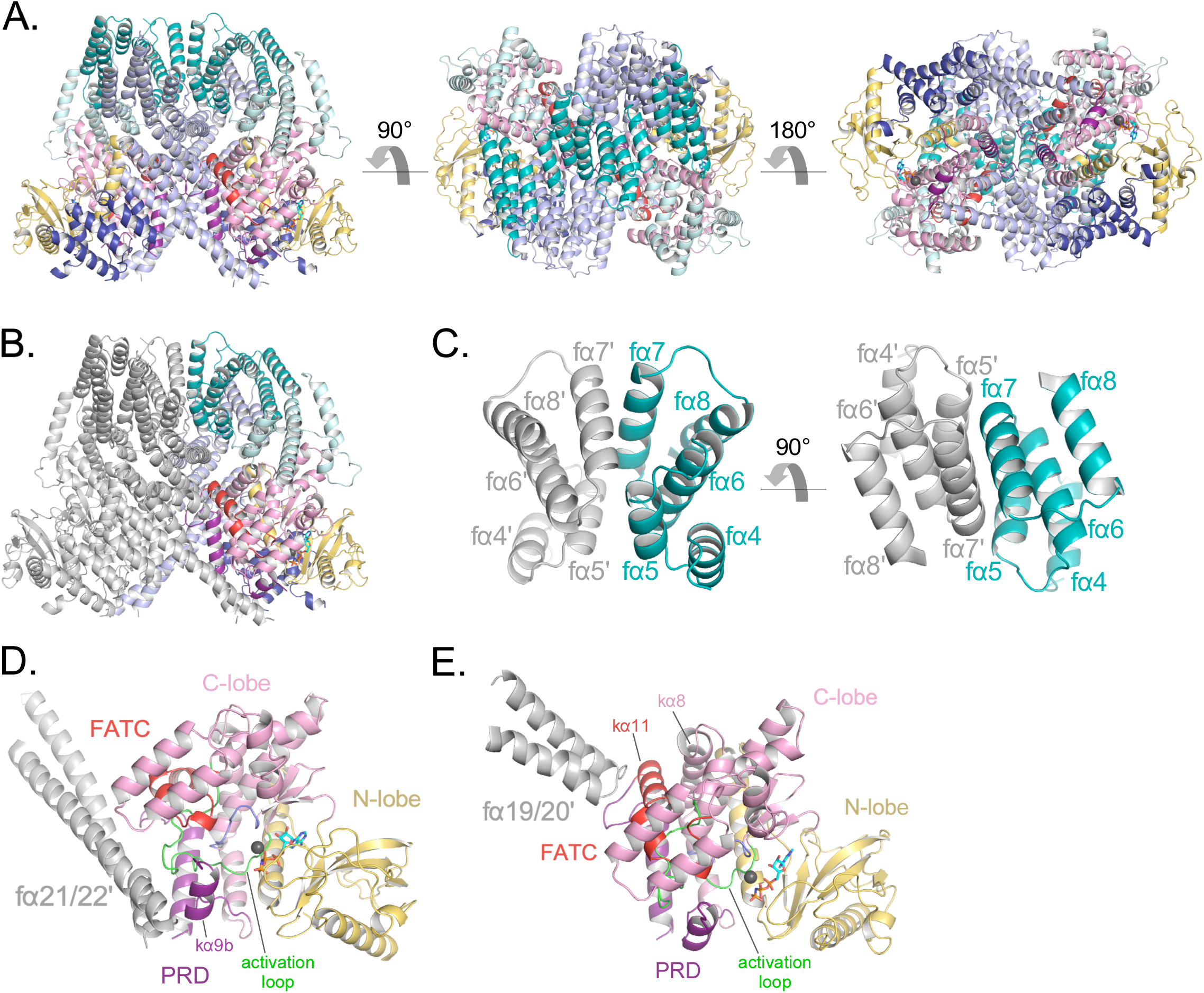
Details of the FAT domain and dimer interface of ATM. **A)** Structure of the FATKD colored by domain as in Figure 1A showing side, top and bottom views. AMP-PNP molecule and single magnesium ion colored cyan and gray, respectively. **B)** Structure of the FATKD with one chain colored gray to highlight intermolecular contacts. **C)** Intermolecular helical packing within the upper dimer interface by FAT helices fα4-8 located within the TRD2 subdomain. **D)** Details of the contacts between the long coiled coil of TRD3 (fα21-22) with the kα9b of the PRD of the symmetric ATM protomer. **E)** Contacts between fα19-20 of TRD3 with the FATC of the symmetric ATM protomer.

TRD2, which is composed of 9 *α* helices (fα4 to fα12), forms a substantial part of the dimer interface, with two separate regions contributing roughly one-third of the surface area buried on dimerization (∼1350 Å^2^ of TRD2 buried per ATM protomer) (Figure 2B). TRD2 helices fα5 to fα8 contact the corresponding TRD2 region of the second protomer in a 2-fold symmetric interface (Figure 2C), while helices fα4 to fα5 contact TRD3 helices fα16 to fα18. The majority of these intermolecular interactions are van der Waals contacts by hydrophobic residues, along with a small hydrogen bond network and electrostatic interactions at the periphery of the hydrophobic residues. Additional buried intermolecular salt bridges between highly conserved residues R2032-E2272 and K2044-E2304 likely function to further stabilize the dimer interface.

TRD3, which is composed of 10 helices (fα13 to fα22), accounts for the largest number of dimerization contacts and for slightly over half the surface area buried. In addition to the aforementioned TRD3-TRD2 intermolecular contacts, TRD3 also contacts the kinase domain of the second protomer. These contacts, which account for approximately one-quarter (∼930 Å^2^) of the surface area buried on dimerization, involve a C lobe surface patch that extends to the edge of the catalytic cleft (Figure 2D and E). The most prominent interactions are made by the TRD3 fα21-fα22 helices, which form a long coiled coil that extends across the dimer interface and packs with the C lobe near the catalytic cleft. These contacts are centered on the C lobe kα9b helix (prefix “k” denotes kinase domain helices) that is part of the PRD (Figure 2D). As reported previously^29, 33–35^, the kα9b helix occludes part of the putative substrate-binding site, and the packing of the coiled coil against it may well stabilize this autoinhibitory conformation. A second set of contacts, made by the TRD3 helices fα19 to fα20, are centered on the FATC kα11 helix located at the end of the C lobe patch, distal from the catalytic cleft (Figure 2E). The FAT domain thus appears to serve multiple functions: it clamps down on the KD N and C lobes in an arrangement thought to be critical in stabilizing the inactive KD conformation of other PIKKs^36^, it is critical for the dimerization of ATM, and it may help stabilize the kα9b conformation that occludes the putative substrate-binding site of the second protomer.

### Kinase domain conformation

The binding of the AMP-PNP cofactor to the catalytic cleft is overall similar those of other PIKKs^29, 30, 36, 37^ (Figure 3A to C). The adenine ring is sandwiched between hydrophobic or aromatic residues from the N lobe (Leu2715, Leu2767 and Trp2769) and C lobe (Leu2877 and Ile2889), while the N6 and N1 groups hydrogen bond to backbone carbonyl and amide groups of Glu2768 and Cys2770, respectively (Figure 3B and C). While the involvement of these residues is conserved among PIKKs, the precise position and orientation of the purine group relative to the C lobe exhibits some variation (up to ∼1.5 Å and ∼20°) among PIKK structures^30, 37^. Similarly, the conformations of the AMP-PNP ribose and phosphate groups in the ATM structure are within the range of variability among PIKK structures, albeit a wider range than that of the purine group. This is possibly due to the use of different ATP analogs, and in the case of mTOR due to conformational change between the inactive and active states^37^.

**Figure 3.**
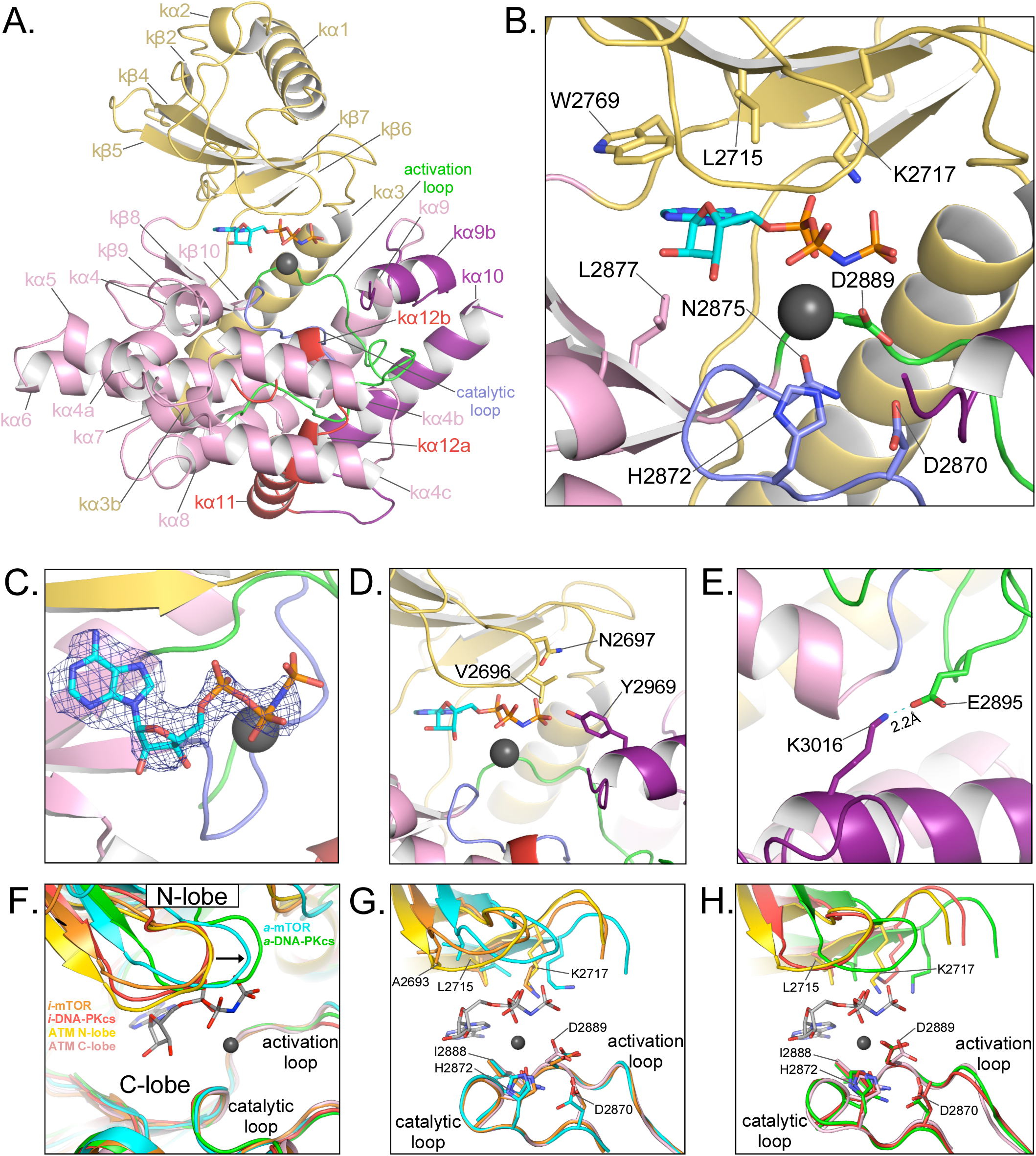
Details of the kinase domain and active site of ATM. **A)** Structure of the Kinase, PRD, and FATC domains with secondary structure elements labeled. Also highlighted are the locations of the catalytic loop (residues 2866-2875, blue) and activation loop (residues 2888-2911, green). AMP-PNP and a single magnesium ion in the active site colored cyan and gray, respectively. **B)** Details of the interactions within the active site. Critical residues within the N lobe, C lobe, catalytic and activation loops are shown as sticks and labeled. **C)** Electron density around AMP-PNP cofactor, contoured to 7σ. **D)** Steric occlusion of the γ phosphate of AMP-PNP by residues V2696 and N2697 on the N lobe and Y2969 on kα9b of the PRD. **E)** Salt-bridge formed between K3016 (PRD, kα10) to E2895 (activation loop). K3016 is acetylated during the DNA-damage response and correlates with active ATM kinase. **F)** Alignment of the Kinase domain of ATM with those of mTOR and DNA-PKcs in the inactive and active states. Structures are aligned along their corresponding catalytic and activation loops. ATM N and C lobes are colored yellow and pink, respectively. Inactive and active mTOR are colored orange and cyan, respectively. Inactive and active DNA-PKcs are colored red and green, respectively. Arrow indicates movement of the N lobe relative to the C lobe coincident with mTOR and DNA-PKcs activation. **G)** Positions of catalytically important residues relative to those of mTOR in the inactive and active states. **H)** Positions of catalytically important residues relative to those of DNA-PKcs in the inactive and active states.

The phosphate groups interact with the N lobe directly, through a contact between Lys2717 and the *α* phosphate group, and with the C lobe indirectly, through the Mg^2+^ cofactor. The position of Lys2717 is equivalent to a critical lysine residue in canonical kinases, where it is thought to orient the *γ* phosphate group for phosphotransfer. Whether the lysine residue has an analogous role in PIKKs is not yet known. In addition to this lysine residue, canonical kinases have an N lobe *β* hairpin, rich in glycine residues, that packs against the phosphate groups when both the ATP and peptide substrates are assembled^46^. The corresponding hairpin in PIKKs is more variable in sequence. The corresponding loop in ATM contains Gly2694 and Gly2695, with the latter’s backbone amide being ∼4 Å away from the *β* phosphate group. This glycine rich loop also contains Val2696 and Asn2697 that, together with Tyr2969 on kα9b, appear to occlude the *γ* phosphate (Figure 3D). The Mg^2+^ ion, which contacts the AMP-PNP *α* phosphate group, is coordinated by Asn2875 and Asp2889 of the C lobe – interactions conserved among both PIKKs and canonical protein kinases^26, 33–36^. Our maps show only a low level of density for a second Mg^2+^ ion thought to be involved in phosphoryl transfer by canonical kinases^47, 48^. The second Mg is often poorly ordered in canonical kinases as well as PIKKs^33–35^.

On the C lobe, the magnesium ligands and other critical catalytic residues map to two loops, named catalytic loop (residues 2866 to 2875) and activation loop (residues 2888 to 2911) by analogy to canonical kinases. As with previous studies of ATM and Tel1, these loops are well ordered and their catalytic residues show clear density in our maps. The catalytic loop contains Asp2870, Asn2875 (a Mg^2+^ ligand) and His2872. Asp2870 acts as the catalytic base to deprotonate and likely orient the hydroxyl group of the incoming substrate^49^ and His2872 is thought to stabilize the transition state of the phosphoryl transfer reaction^36^. The activation loop is named for its conformational change in the activation of canonical kinases, where it makes up part of the polypeptide substrate-binding site. Previous studies showed that the activation loop of ATM/Tel1 packs with the k*α*9b helix of the PRD, and by analogy to canonical kinases it was suggested that the k*α*9b helix may block substrate binding. This was recently confirmed by the cryo-EM analysis of the PIKK Smg1, which, like ATM, is specific for a glutamine residue in the position (P_+1_) after the serine/threonine phosphorylation site. In the Smg1-UPF1 substrate structure^30^, the glutamine side chain of the peptide substrate makes a pair of hydrogen bonds to backbone amide and carbonyl groups on the activation loop. This interaction is effectively mimicked in our ATM structure by the side chain of Gln2971 on the k*α*9b helix of ATM (Supplemental Figure 8).

It has been suggested that the Tel1/ATM active site residues are in a catalytically competent conformation, and thus the occlusion of the substrate-binding site would the main mechanism of keeping the kinase autoinhibited^29, 33–35^. However, studies of mTOR have shown its activation involves a conformational change in the FAT domain, which in turn allows the KD N and C lobes to move relative to each other. This results in the realignment of catalytic residues on the N and C lobes relative to each other, bringing them into the correct register for efficient catalysis^37^. The recently reported active-state structure of DNA-PKcs recapitulates the N-C lobe conformational change on activation^39^, raising the possibility that this is a common activation mechanism of PIKKs, irrespective of their distinct activators and N-terminal solenoid structures to which these activators bind.

Thus, to help evaluate whether the ATM active site residues are in a catalytically competent conformation, we superimposed the KD domains of ATM, mTOR and DNA-PKcs, the latter two in both their active and inactive conformations, by aligning their C lobe catalytic and activation loops. In this superposition (Figure 3F), the ATM N lobe is positioned remarkably similar to those of inactive mTOR and inactive DNA-PKcs, with the three N lobes forming a tightly clustered set clearly distinct from the cluster of active mTOR and active DNA-PKcs N lobes. The positions of the N lobe hairpin and other ATP-interacting residues relative to the C lobe catalytic and magnesium-coordinating residues are much closer to those of the inactive mTOR and DNA-PKcs structures compared the active ones (Figure 3G and H). This positioning thus suggests that in addition to relieving the blockage of the substrate-binding site, the activation of ATM may well involve a conformational change within the FAT domain and the associated change in the relative orientation of the N and C lobes of the kinase domain.

### Structure of ATM bound to the C-terminus of Nbs1

Previous studies indicated that the Xrs2 C-terminal FxF/Y motif and the acidic region that immediately precedes it bind to two Tel1 regions spanning the Spiral and Pincer domains near the elbow^22, 35^ (Figure 4A). We thus made cryo-EM grids of human ATM (1.2 μM) mixed with a peptide (207 μM) that corresponds to the C-terminal 28 residues of human Nbs1 and which encompasses the acidic region and FxF/Y motif (residues 727-754, hereafter referred to as Nc28).

**Figure 4.**
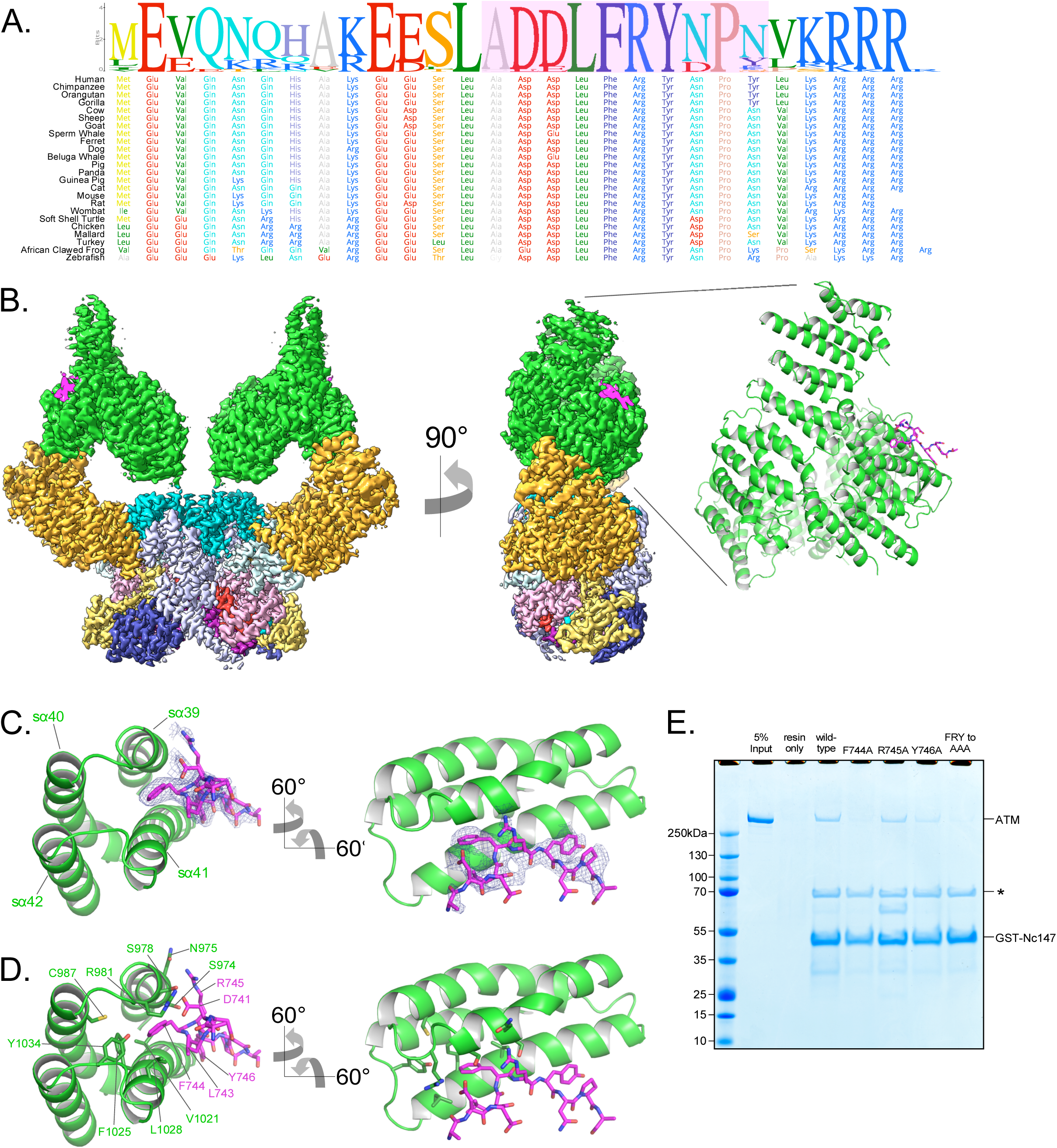
Structure of ATM bound to the C-terminus of Nbs1. **A)** Alignment of the Nbs1 C-terminal 28 residues (Nc28) from 24 vertebrate species showing the conservation of the C-terminus and FxF/Y motif. Portion of Nc28 visible in the cryo-EM map is highlighted in magenta. **B)** Location of the Nc28 bound to the ATM Spiral domain in the composite map (left) and structure (right). Nc28 is colored magenta. **C)** Zoom of the location of the Nc28 peptide showing Phe744 inserted into a hydrophobic groove made up of ATM helices sα39 to sα42. Nc28 peptide density is contoured to 5σ. **D)** Details of the interaction between Nc28 and the ATM Spiral, showing Phe744 inserted into a hydrophobic pocket created by ATM residues Arg981, Cys987, Val1021, Ala1024, Phe1025, Leu1028, and Tyr1034. Arg745 of Nbs1 makes electrostatic contacts with ATM Asn975 and Ser978. Tyr746 of Nbs1 packs against the sα41. Nbs1 Leu743 makes contacts with an adjacent hydrophobic cleft created by ATM residues Leu1028 and His1064. Nbs1 Asp741 makes electrostatic and H-bonding contacts with ATM residues Arg981 and Ser978, respectively. **E)** ATM pull-down assay using GST-tagged Nbs1 C-terminal 147 residues (GST-Nc147) as bait. Lanes are labeled at the top of the image, and bands are labeled on the right. Asterisk indicates Hsp70/DnaK contaminant that co-purifies with all GST-Nc147 preparations.

The cryo-EM data yielded a consensus reconstruction in point group C2 that extended to 2.6 Å resolution as determined from the gold standard FSC (Supplemental Figure 9). The initial map had additional density, absent in the apo ATM (Supplemental Figure 9D), at the ATM Spiral domain. Partial signal subtraction and symmetry expansion procedures followed by iterative focused 3D classifications identified 75 % of the particles that had the extra density (Supplemental Figure 3A and 9C). After focused refinements of the ATM Spiral domain, we built a 10-residue Nbs1 segment (^740^ADDLFRYNPY^749^) into the improved density (Figure 4B). There is no interpretable density for residues 727 to 739 preceding the built segment or residues 750 to 754 after. We note that in contrast to previous yeast two-hybrid assays indicating that the C-terminus of Xrs2 interacts with both the Spiral and Pincer domains of Tel1^22^, we find no evidence of the ATM Pincer domain being involved in binding to the Nbs1 peptide.

The Nbs1 peptide adopts an overall extended conformation except for one turn of a 3_10_ helix in the middle (residues Asp742 to Arg745; Figure 4C and D). It binds to a hydrophobic groove between two helical repeats formed by helices sα39 to sα42 (denoted “s” for Spiral domain helices). The most extensive ATM contacts are made by Nbs1 Leu743 and Phe744, which pack together preceding the 3_10_ helix. Phe744 inserts deepest into the hydrophobic groove and makes van der Waals contacts to the side chains of Ser978, Arg981, Cys987, Val1021, Ala1024, Phe1025, Leu1028, and Tyr1034 of ATM (Figure 4C and D). Leu743, which is closer to the solvent exposed surface of the ATM groove, packs with Ala1024, Leu1028 and His1027 (Figure 4C and D). Of the two acidic residues that precede the Leu743-Phe744 pair, Asp741 hydrogen bonds with Ser978 and Arg981, while Asp742 is solvent exposed and uninvolved in ATM contacts (Figure 4C and D). Arg745, the last residue of the 3_10_ helix, has weaker side chain density, although its guanidinum group is positioned between the side chains of Asn975 and Ser978 (Figure 4C and D). Tyr746 is the last Nbs1 residue that contacts ATM. Its side chain, which is stabilized by intramolecular stacking with Pro748, packs with side chain and backbone groups of Gly1016, Gln1017 and Thr1020 of ATM (Figure 4C and D).

The ATM residues that make up the Nbs1 binding site, and in particular those at the Leu743-Phe744 binding pocket, are highly conserved compared to the rest of Spiral domain residues (Supplemental Figure 4 and Supplemental Figure 10A). In addition, many of these ATM residues have been found mutated in various cancers (Supplemental Figure 5D). Ser978 is a hotspot for cancer-associated missense mutations that are predicted to disrupt this interaction (S978P, S978A, S978Y, Supplemental Figure 5D and E). Other cancer-associated missense mutations at the Nbs1 binding groove occur at lower frequencies. The structure predicts that these mutations, which include S974F, R981C/H, C987Y/W, and F1025S/L, are also likely to disrupt the ATM-Nbs1 interaction. The abundance of these mutations may indicate that the ATM-Nc28 interaction is functionally important for the ATM-mediated DNA damage response. Reported mutations in Nbs1 are fewer, but include the F744L mutation that would disrupt the most critical portion of the ATM-Nc28 interaction (Supplemental Figure 5F).

The structural elements of ATM including the hydrophobic pocket and adjacent cleft are nearly identical in both the apo and Nbs1-bound structures, indicating that Nbs1 binding does not induce any noticeable structural rearrangements to this region (Supplemental Figure 9D). Additionally, no significant changes to the structure of the kinase domain were observed when comparing the unbound and Nc28-bound states (Supplemental Figure 10D). Consistent with these observations and with reports that ATM activation requires all 3 members of the MRN complex and dsDNA^7^, we observed no increase in ATM catalytic activity in steady-state kinase assays containing high concentrations of this peptide (Supplemental Figure 10C). Finally, though previous studies suggest that the Nbs1 C-terminus may interact with the ATM FATC domain^33, 50^, we observe no additional electron density within this region of ATM despite our sample containing a large molar excess of the Nc28 peptide.

The N-terminal acidic half of the peptide is not visible in our map. However, the orientation of the visible portion indicates that it likely extends into a basic patch created by ATM residues Arg981, Arg982, and Lys1033, raising the possibility that the acidic residue clusters (Glu728, Glu736 and Glu737) contribute to binding through long-range electrostatic interactions (Supplemental Figure 10B). Similarly, while the C-terminal 4 residues of Nbs1 (^750^LKRRR^754^) are also not visible, they would be in the vicinity of an acidic patch of ATM (Glu964, Asp965, Asp1007, Glu1009, and Asp1013; Supplemental Figure 10B).

To evaluate the relative contributions of the aforementioned contacts, we made several mutations in the Nbs1 FxF/Y motif and tested the ability of these mutant peptides to bind to ATM in a GST pull-down assay. Four mutations (F744A, R745A, Y746A, and ^744^FRY^746^ to ^744^AAA^746^) were introduced into a GST-tagged Nbs1 C-terminal 147 residue polypeptide (GST-Nc147). While the wild-type GST-Nc147 and the R745A mutant both enriched for ATM in pull-down assays, all other mutations partially disrupted this interaction (Figure 4E). F744A and FRY to AAA mutations appeared to disrupt the interaction almost completely, whereas Y746A showed an intermediate effect. These results indicate that Nbs1 Phe744 is central to ATM binding, Tyr746 clearly contributes, while Arg745 makes at most a minor contribution compared to the other two residues (Figure 4C and D).

The Mre11 binding motif of Nbs1 (residues 682-693) is located approximately 60 residues N-terminal to the FxF/Y motif, indicating that the MRN complex likely binds along the Spiral and/or Pincer domains of ATM. Interestingly, we mapped multiple patches of conserved residues to the inner portion of the Spiral domain α-solenoid, which may represent binding sites for MRN-dsDNA or other effector proteins (Supplemental Figure 7B). Notably, the Nbs1 interaction site is located over 100 Å away from the active site. This is reminiscent of mTOR and DNA-PKcs, whose activators, respectively Rheb and Ku-DNA, bind far from the KD and activate their respective kinases allosterically^37, 39^. This mechanism would suggest that the full MRN-dsDNA complex induces structural changes that are transmitted through the Spiral and Pincer domains to the FATKD, although the Rad50 coiled coil domain may be long enough to extend from the head of the MRN complex and make direct contacts to the kinase domain and/or FATC to promote activation.

### Biochemical analysis of MRN-mediated ATM activation and the role of the Nbs1 C-terminus

To better understand how human ATM kinase is activated and to further evaluate the role of the Nbs1 C-terminus, we expressed the human MRN complex in mammalian cells via transfection of a polycistronic vector (see Materials and Methods) and purified it to near homogeneity (Supplemental Figure 11A). Each protein contains a C-terminal FLAG tag and Mre11 harbors the H129N mutation, which abolishes nuclease activity without interfering with DNA binding and ATM activation^51^. We first evaluated the effect of increasing concentrations of the purified MRN complex on the phosphorylation of a p53 substrate peptide by ATM. In keeping with findings with the Tel1-MRX homologs^23^, MRN alone stimulated the ATM kinase activity only modestly, by a factor of ∼4 (Supplemental Figure 11B). The half-maximal effective concentration (EC_50_) of MRN was ∼27 nM, indicative of a high affinity for ATM.

We next tested linear dsDNA fragments of lengths ranging from 100 to 2000 base pairs (bp) for their ability to stimulate ATM (25 nM) in the presence of MRN (250 nM; concentration ∼10-fold above its EC_50_ towards ATM alone). The dsDNA fragments were added at a constant mass concentration of 2.5 ng/µL (nucleotide concentration of 3.85 µM base pairs). dsDNA lengths greater than ∼200 bp lead to maximal ATM activation, representing an additional ∼25-fold increase in p53 phosphorylation relative to ATM-MRN alone, or a ∼100-fold increase relative to ATM alone (Figure 5A, Supplemental Figure 11F). This dsDNA length dependency for ATM-MRN activation is qualitatively similar to that reported for the Tel1-MRX complex^23^. We then performed a dose response analysis for dsDNA of 100, 250 and 500 bps lengths. As shown in Figure 5B, 250 and 500 bp dsDNA stimulated with very similar EC50 values of 1.8 nM and 1.5 nM, while stimulation by 100 bp dsDNA was a factor of 5 lower, compared to the maximal levels of the longer DNA fragments, even at the ∼1000-fold higher concentration of 2.3 µM (Figure 5B and Supplemental Figure 11C). This suggests that the low level activation of shorter dsDNA fragments is likely not due to their reduced affinity for the ATM-MRN, and that structural aspects of longer DNA, such as the ability to link distal binding sites on ATM-MRN, may be important. Stimulation by dsDNA is strictly dependent on the presence of the MRN complex, as in the absence of MRN, 350 bp dsDNA at a concentration of 25 nM, which is more than 10-fold above the EC_50_ of the 250 and 500 bp DNA failed to activate (Supplementary Figure 11D and F).

**Figure 5.**
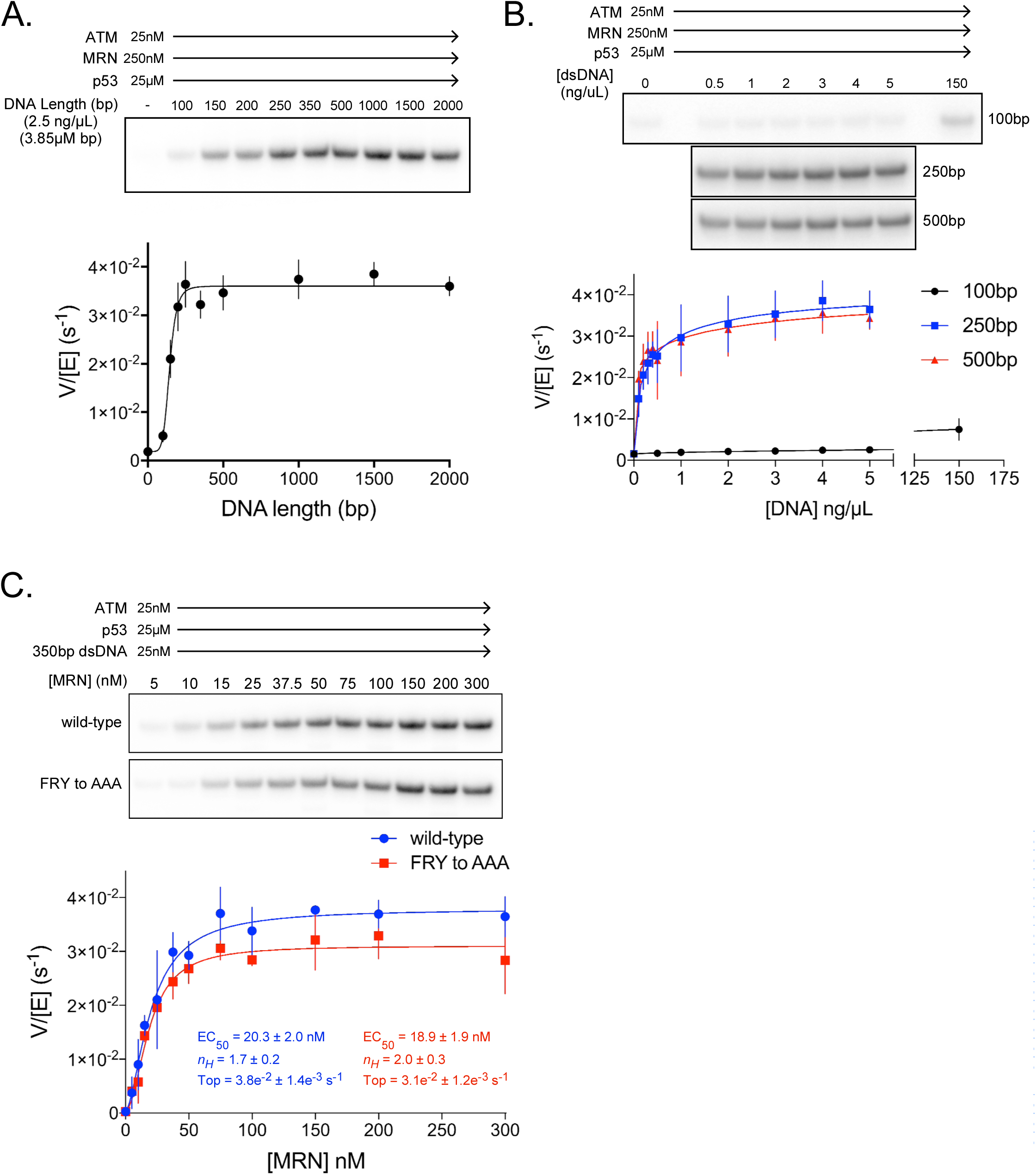
ATM activation requires MRN and long dsDNA, but not the FxF/Y motif of Nbs1. **A)** <u>Top</u>: Steady-state ATM kinase assay using 25 nM ATM, 250 nM MRN and 25 µM p53 substrate with 2.5 ng/µL of dsDNA of various lengths added. <u>Bottom</u>: Quantification of ATM enzymatic velocity as a function of DNA length. 3 biological replicates per point. Curve fit to the EC_50_ equation. **B)** <u>Top</u>: Steady-state ATM kinase assay using 25 nM ATM, 250 nM MRN and 25 µM p53 substrate with various concentrations (0.5 to 5 ng/µL) of 100bp, 250bp and 500bp dsDNA fragments. 150 ng/µL 100bp dsDNA is in the upper right lane. <u>Bottom</u>: Quantification of ATM enzymatic velocity as a function of DNA concentration. 3 biological replicates per point, Curve fit to the EC_50_ equation. **C)** <u>Top</u>: Steady-state ATM kinase assay using 25 nM ATM, 25 nM 350bp dsDNA, and various concentrations of wild-type or FRY to AAA mutant MRN. <u>Bottom</u>: Quantification of ATM enzymatic velocity as a function of MRN concentration. 3-6 biological replicates per point. Curves fit to the EC_50_ equation.

We next evaluated whether MRN-dsDNA affects the intrinsic catalytic step or peptide substrate binding. The steady state kinetic analysis of ATM phosphorylating the p53 substrate showed that MRN-dsDNA increased the catalytic step 460-fold, with *k*_cat_ values of 0.0005 s^-1^ and 0.24 s^-1^ in the absence and presence of MRN-DNA, respectively, while the *K*_M_ values were unchanged within experimental error (58 µM and 67 µM; Supplemental Figure 11D and E). Time course assays performed under single-turnover conditions also indicate that MRN directly increases the rate of the catalytic step, as opposed to the hypothetically rate limiting product release step (Supplemental Figure 11G).

We next made MRN containing the Nbs1 ^744^FRY^746^ to ^744^AAA^746^ mutation and titrated this mutant complex into kinase assays with saturating amounts of 350bp dsDNA present (Figure 5C). In contrast to pull-down assays, we observe only a minor defect in MRN-mediated activation of ATM kinase activity with this mutant. The maximal catalytic rate of ATM with this mutant was approximately 18 % lower compared to the wild-type MRN complex, although an effect on the EC_50_ of activation was not discernible. It is likely that the interactions of ATM with Mre11 and Rad50, and possibly with DNA largely compensate for the weaker binding of the Nbs1 mutant under our *in vitro* assay conditions.

## Discussion

Our cryo-EM structure of human ATM shows an overall symmetric dimer, albeit with the Pincer and Spiral domains exhibiting some mobility arising from conformational flexibility within the Pincer domain. Nevertheless, this conformational flexibility does not translate to any noticeable structural rearrangements within the FATKD segment, nor does it break the inherent symmetry of this segment (Supplemental Figure 6). Previous cryo-EM studies of human ATM and yeast Tel1 have observed asymmetric dimers and monomers, the latter of which was shown to have increased catalytic activity^29, 32^. In our purifications and subsequent cryo-EM structure determination, we failed to find any particles representing asymmetric dimers or monomers, also consistent with the most recent cryo-EM structures of Tel1^34, 35^. While it is not clear how the asymmetric dimers and monomers arise, it is conceivable that expression and purification conditions may play a role. Detergents have previously been demonstrated to disrupt the ATM dimeric structure and increase ATM kinase activity independent of MRN^52^. We note that the report of active monomer preparations demonstrated only an approximately 10-fold increase in catalytic activity relative to the inactive dimeric state^32^, whereas our quantitative kinase assays show a ∼100-fold activation upon MRN+long dsDNA addition (Supplemental Figure 11F).

As noted in previous ATM and Tel1 structures^29, 33–35^, the apparent reason for the inhibited state of the ATM dimer is the k*α*9b helix packing against the putative substrate-binding site and inserting a glutamine side chain into the activation loop, at a site where a glutamine side chain from the substrate would bind (Supplemental Figure 8)^30^. As the k*α*9b helix packs with the TRD3 coiled coil of the second protomer in the dimer, an interaction that possibly stabilizes the k*α*9b conformation, it was proposed to couple ATM dimerization to autoinhibition^29, 33–35^. In the Smg1 cryo-EM structure, where Smg1 is in complex with the Smg8 and Smg9 subunits and is thought to be in an active state, the entire ka9b helix is disordered^30^.

The k*α*9b helix is part of the PRD domain, which was first described based on a yeast two-hybrid screen to identify the part of the ATR PIKK that interacts with its activator TopBP1^53^. This identified a minimal ATR fragment that encompasses k*α*9b and k*α*10, although the latter is an integral and unchanging part of the PIKK C lobe structure and likely not involved in regulating the kinase. In ATM, the k*α*9b and k*α*10 helices are connected by 27-residue loop (residues 2975-3000) that is not conserved across species and is not visible in our map (Supplemental Figure 4). An unstructured loop intervening the k*α*9b and k*α*10 helices is present in most PIKK structures reported to date, including a >1000 residue insertion in Smg1^30^.

In structures of inactive DNA-PKcs the k*α*9b and subsequent k*α*9c (unique to DNA-PKcs) helices also pack against the substrate-binding site and occlude it^38, 39^. However, even though DNA-PKcs is glutamine directed as ATM, it does not have the equivalent of the Gln2971-activation loop interaction of ATM. Nevertheless, upon DNA-PKcs activation, the pair of k*α*9b and k*α*9c helices move to an alternate packing position on the C lobe and fully expose the substrate-binding site^39^. The movement of k*α*9b-k*α*9c is likely due entirely to the realignment of the N and C lobes of DNA-PKcs, as this segment only interacts with the kinase domain (DNA-PKcs does not have the equivalent of the ATM TRD3 coiled coil packing with and possibly stabilizing k*α*9b across a dimer interface). By contrast, in mTOR, which is not glutamine directed and instead prefers a hydrophobic residue at the P_+1_ position, the k*α*9b helix does not block the putative substrate-binding site, nor does it change conformation on activation^37^. Thus, while k*α*9b may play a role in ATM autoinhibition, this is not a conserved feature of the PIKK family.

Rather, several lines of evidence point to the realignment of the N and C lobes of the kinase domain as the key event in the activation of ATM. As discussed above, the relative orientation of the N and C lobes of ATM is remarkably similar to those of the inactive-state mTOR and DNA-PKcs structures, and distinct from those of the active-state counterparts, which cluster together (Figure 3F). The proper alignment of the N and C lobes is central to kinase activation, as they contain residues critical for substrate binding and catalysis^37^.

The mechanism of a PIKK activation was initially established by comparing the inactive and active states of the mTOR holoenzyme^37^. This work pointed to the FAT domain being the key autoinhibitory element that keeps the N and C lobes and their active site residues in an unproductive configuration^37^. Binding of the mTOR activator, the small GTPase Rheb, causes a motion of the N-heat solenoid (comparable to the ATM Spiral in its location in the primary sequence but not in its structure). The motion of the N-heat solenoid pulls and twists the FAT domain on which it is anchored, causing it to undergo an extensive conformational change. This shifts the HRD of the FAT domain away from the N lobe, allowing the N lobe to relax to a productive configuration relative to the C lobe. The inhibitory effect of the FAT domain is supported by activating mTOR mutations, which map to residues that couple the N and C lobes and the FAT domain^37^.

The recent cryo-EM structures of activated DNA-PKcs recapitulate the FAT conformational change observed in mTOR^39^. Binding of the Ku70-Ku80-dsDNA complex causes the DNA-PKcs N-heat solenoid (analogous to ATM Spiral and mTOR N-heat) anchored on the FAT domain to move, allosterically inducing a conformational change in the FAT domain that results in realigning the N and C lobes. In both mTOR and DNA-PKcs, the N-heat solenoids that move are anchored between the FAT TRD2 and TRD3 domains, at essentially the same location. Furthermore, even though their overall N-heat solenoids are structurally unrelated, they both use a four-helix bundle followed by a loop to interact with their respective FAT domains. Our ATM structure reveals a remarkably similar interface between the FAT domain and the Pincer (equivalent to M-heat of mTOR and DNA-PKcs) from same protomer. The similarities include an identical TRD2-TRD3 site of the FAT domain and a four-helix bundle and loop element from the Pincer that binds to it, in essentially the same configuration as those of mTOR and DNA-PKcs (Supplemental Figure 12). Based on this structural conservation and the high level of FAT conservation across the PIKK family, it is conceivable that ATM activation too involves this same mechanistic step, with the activator MRN-dsDNA complex triggering the Pincer to pull and twist the FAT domain to realign the N and C lobes.

Because the FAT domain plays a central role in the dimerization interface of the inactive ATM dimer, a conformational change in the FAT domain may well affect the relative arrangement of the two ATM protomers. This may disrupt the packing of k*α*9b with the TRD3 coiled coil from the second protomer, allowing the movement of k*α*9b to expose the P_+1_ substrate-binding site. Our steady state kinetic analysis showed that MRN+dsDNA increases the *k*_cat_ value of ATM phosphorylating a p53 substrate by over two orders of magnitude without affecting the *K*_M_ value (Supplemental Figure 11D to F). This mirrors the steady-state kinetic constants of mTOR activation^37^. However, activation having no effect on the *K*_M_ value was unexpected, given that the presumed binding site for the P_+1_ position of the substrate is blocked in the structure. This may be analogous to findings with the canonical Cdk2-CyclinA kinase, which is activated by phosphorylation on its activation loop. Even though phosphorylation reorganizes the substrate-binding site on the Cdk2 activation loop, the majority of the increase in catalytic efficiency is reflected in an increased rate of phosphoryl group transfer step^54^. Based on the model proposed for Cdk2^54^, it is possible that the blocked P_+1_ position in inactive ATM allows substrate binding, but with the phospho-acceptor group in an unfavorable position or orientation that reduces the rate of phosphoryl group transfer.

In conclusion, our structural data supports the model that activation of the ATM kinase domain may be mechanistically analogous to the activation of mTOR and DNA-PKcs. While it is not known how Rad50 and Mre11 interact with ATM, we find that the Nbs1 FxF/Y motif binds to the ATM Spiral domain, and the Spiral domain of Tel1 was shown to bind to dsDNA independently of MRN^34^. Thus, it is possible that the Spiral domain is involved in triggering a motion of the Pincer domain. This would then result in a conformational change in the FAT domain that not only realigns the N and C lobes of the kinase domain but also relieves the blockage of the substrate-binding site.

## Methods

### Peptides, reagents, and antibodies

A peptide corresponding to the C-terminal 28aa of Nbs1 (Nc28) was purchased from Genscript and dissolved in 100mM HEPES pH 7.4 to a concentration of 2mM as measured by 205nm absorbance. No other specialty reagents or antibodies were used in this study.

### Purification of ATM kinase

Codon optimized human ATM harboring a N-terminal FLAG tag was purified from a stably transfected HEK 293 cell line grown in suspension. Cells were grown to a density of 2-3e^6^ per liter, pelleted, and resuspended in 50mL per liter of ATM lysis buffer containing 50mM Tris-HCl pH 8.0, 500mM NaCl, 50mM KCl, 1mM EDTA, 10% glycerol, 0.5mM TCEP, supplemented with protease inhibitors aprotinin, leupeptin, pepstatin, AEBSF. Cells were lysed by 2 passages through a cell disruptor, and cell lysate was clarified by centrifugation. Soluble cell lysate was incubated with α-FLAG M2 sepharose (Sigma) for 1 hour at 4°C. Lysate was passed over a gravity column, and resin was extensively washed with lysis buffer. FLAG-ATM was eluted from the resin in lysis buffer containing 0.2 mg/mL FLAG peptide. FLAG-ATM was diluted to ∼100mM Na/KCl and loaded onto a Mono-Q 5/50GL column equilibrated in Buffer A (25mM Tris pH 8.0, 100mM NaCl, and 0.5mM TCEP). ATM was eluted from the column with a linear gradient of 0-100% buffer B (25mM Tris pH 8.0, 1M NaCl, and 0.5mM TCEP). ATM eluted at approximately 325mM NaCl. Peak fractions were pooled and used directly for cryo-EM grid preparation. The remaining fractions containing ATM were pooled and glycerol was added to a final concentration of 10%. ATM was aliquoted, flash frozen in liquid N_2_, and stored at -80°C for kinase and pull-down assays.

### Purification of the MRN complex

Codon optimized human Mre11, Rad50, and Nbs1 each containing their own CMV promoters, non-cleavable C-terminal FLAG tag, and poly-A sequences were cloned into a polycistronic mammalian expression vector. Mre11 also harbored the H129N mutation that abolished nuclease activity. FLAG-tagged MRN was expressed by PEI transient transfection of HEK 293 cells growing in suspension at a density of 1-2e^6^ per liter. After 48 hours cells were pelleted and resuspended in 50mL per liter of MRN lysis buffer containing 50mM Tris-pH 8.0, 750mM NaCl, 50mM KCl, 10% glycerol, 0.5mM TCEP, supplemented with protease inhibitors aprotinin, leupeptin, pepstatin, AEBSF. The remainder of the purification procedure was performed using the same protocol as ATM. Peak Mono-Q fractions were pooled, aliquoted, flash frozen in liquid N_2_, and stored at -80°C for kinase assays. The MRN FRY-to-AAA mutant was generated using InFusion mutagenesis and purified in the same manner as wild-type MRN.

### Cryo-EM grid preparation

For the apo ATM sample, Mono-Q purified ATM kinase was diluted to 0.6 mg/mL in 25mM Tris pH 8.0, 300mM NaCl, 0.5mM TCEP, 1.25mM AMP-PNP, and 2.5mM MgCl_2_ and incubated on ice for 1 hour. For the ATM-Nc28 sample, frozen ATM was thawed and dialyzed against 25mM Tris pH 8.0, 250mM NaCl, and 0.5mM TCEP overnight. ATM was then mixed with the Nc28 peptide such that the final concentrations of each were 1.2µM (0.42 mg/mL) and 207µM (0.73 mg/mL), respectively. 2.5mM AMP-PNP, and 5mM MgCl_2_ were spiked in and the sample was incubated on ice for 1 hour. Each sample was briefly centrifuged to remove large aggregated species. 2.5+2.5µL of the samples were applied to glow-discharged UltraAuFoil 300 mesh R1.2/1.3 grids (Quantifoil) via double-sided application. Grids were blotted for 1.5 to 3 seconds at 22°C and 95% relative humidity and plunge frozen in liquid ethane using a FEI Vitrobot Mark IV.

### Cryo-EM data collection

For the apo ATM sample, a single 3-day dataset was collected at the Memorial Sloan Kettering cryoEM Facility on a Titan Krios microscope operated at 300kEV equipped with a Gatan K3 Summit direct electron detector. Data was acquired using a defocus range of - 0.6 to -1.6µm and a pixel size of 1.056Å. Each micrograph was acquired using a 3 second exposure and fractionated into 40 frames with a dose of 20 electrons per pixel per second. A total of 9,028 micrographs were collected using 3X3 image shift. For the ATM-Nc28 sample, a single 3 day dataset was collected at Janelia Research Campus on a Titan Krios microscope operated at 300kEV equipped with a Gatan K3 Summit direct electron detector and an energy filter. Data was acquired using a defocus range of -1.0 to -2.5µm and a pixel size of 1.078Å. Each micrograph was acquired using a 3 second exposure and fractionated into 40 frames with a dose of 20 electrons per pixel per second. A total of 7,866 micrographs were collected using 3X3 image shift.

### Data processing and structure refinement

Beam-induced sample motions were corrected using MotionCorr2 software, and contrast transfer function parameters were estimated using CTFFIND4 software. All subsequent processing steps were performed using RELION-3 software. All reported resolutions are calculated from gold-standard refinement procedures with the FSC=0.143 criterion after post-processing by applying a soft mask, correction for the modulation transfer function (MTF) of the detector, temperature-factor sharpening, and correction of FSC curves to account for the effects of the soft mask as implemented in RELION. **<u>For unbound ATM</u>**, all processing steps are summarized in Supplemental Figures 2 and 3. 8,973 micrographs were selected based on a CTF estimated resolution cutoff of 10Å. ∼2.5 million particles were autopicked using 2D templates generated from a previously collected, smaller ATM dataset collected using a Gatan K2 Summit direct electron detector. Picked particles were cleaned by multiple rounds of binned and unbinned 2D and 3D classifications. The remaining ∼1.5 million particles were subjected to 2 rounds of per-particle Bayesian training, polishing and CTF refinement taking per particle astigmatism and beam tilt into account. Additional rounds of 2D and 3D classifications were performed after polishing to remove truncated ATM dimers. The resulting 303,604 particles were refined while applying C2 symmetry to an overall resolution of 2.50Å. An additional round of 3D classification using 6 classes was performed to subclassify the overall ATM dimer particles into open and closed states. The most open class containing 44,152 particles and the most closed class containing 44,114 particles were each refined while applying C2 symmetry to an overall resolution of 2.75Å and 2.78Å, respectively. For each particle set (overall, open, and closed), partial signal subtraction and symmetry expansion procedures were performed to align the ATM protomer using a soft mask generated from an initial build of the ATM protomer^42^. For each particle set, refinements were performed using masks generated for the overall promoter, HEAT domain (roughly corresponding to residues 1-1588), and FAT+Kinase domains (roughly corresponding to residues 1588-3056). An initial model was docked into each focused map in Chimera software, and focused maps were combined using the combine_focused_maps tool in Phenix software with the local_weighting flag set to True and local_residues set to 10. The final overall, open, and closed protomer structures were iteratively built and refined against these composite maps using Coot and Phenix software. All structure and model-to-map validations were performed using Molprobity tools as implemented in Phenix software using composite protomer maps. Final overall, open, and closed dimer structures were generated by applying NCS operators calculated from the corresponding C2 symmetric dimer maps and refined into the corresponding dimer composite maps. **<u>For the data set of the ATM-Nc28 complex</u>**, all processing steps are summarized in Supplemental Figure 9. 6,531 micrographs were selected based on a CTF estimated resolution cutoff of 5 Å. ∼2 million particles were autopicked using 2D templates generated from a previous ATM dataset collected using a Gatan K2 Summit direct electron detector. Picked particles were cleaned by multiple rounds of binned and unbinned 2D and 3D classifications, with truncated ATM dimers removed prior to polishing. The remaining 287,615 particles were subjected to 2 rounds of per-particle Bayesian training, polishing and CTF refinement also taking per particle astigmatism and beam tilt into account. The resulting 287,615 polished particles were refined while applying C2 symmetry to an overall resolution of 2.52 Å. Partial signal subtraction, symmetry expansion, and focused refinement procedures were performed in the same manner as unbound ATM. Focused maps were combined in the same manner as unbound ATM, and an initial model was refined against the composite map. At this point in the model building we identified extra density adjacent to a hydrophobic cleft in the Spiral domain of ATM. At this stage, a 7-residue segment (^742^DLFRYNP^748^ ) of the Nbs1 peptide had unambiguous density and was built into the map. We then used this model to generate a soft mask around this area including ATM residues 917 to 1101. We next performed iterative rounds of alignment-free focused 3D classification in RELION using particles pre-aligned in the Spiral domain, 3 classes, the skip_align flag, and tau_fudge factor set between 150 to 600 to separate bound and unbound particles. After multiple rounds of 3D local classification, one class containing ∼75 % of the original particles had clear density within the pocket. Focused refinements were repeated and maps were again combined using this subset of particles. The peptide density was better defined in the resulting composite map and we were able to extended the Nbs1 model to 10-residues (^740^ADDLFRYNPY^749^). The final protomer was iteratively built and refined against this composite protomer map using Coot and Phenix software. All structure and model-to-map validations were performed using Molprobity tools as implemented in Phenix software. The final ATM-Nc28 dimer was generated by applying NCS operators calculated from the C2 symmetric dimer map and refined into the dimer composite map.

### Pull-down assays

GST-tagged Nbs1 C-terminal 147 residues (GST-Nc147) harboring a C-terminal StrepII tag was expressed from a pGEX vector in *E. coli* BL21[DE3] cells. Cells were lysed by sonication and protein was initially purified by glutathione sepharose column. Elutions were further purified by StrepTactin Sepharose column and eluted with 10mM desthiobiotin (DTB). GST-Nc147 and mutants were concentrated and DTB was removed by serial buffer exchanges. Proteins were aliquoted, flash frozen in liquid N_2_, and stored at -80°C. For pull-down assays 3µg of GST-Nc147 wild-type and mutants were coupled to 25µL of pre-equilibrated glutathione sepharose resin in 200µL of binding/wash (B/W) buffer containing: 25mM Tris pH 8.0, 200mM NaCl, 0.5mM TCEP, and 0.01% NP-40 for 1 hour rotating at 4°C. Resin was washed 1X 500µL of B/W buffer, and resuspended in 200µL of B/W buffer containing 0.05 mg/mL of FLAG-ATM (10µg per reaction) and incubated for 2 hours rotating at 4°C. The resin was washed 3X with 500µL of B/W buffer for 10 minutes rotating at 4°C. 25µL of pelleted resin was resuspended in 25µL of 2X LDS sample buffer and boiled for 10 minutes. 25µL (50% of the reaction) was loaded onto a 4-12% Bis-Tris SDS-PAGE gel. Proteins were separated by electrophoresis and gels were stained in InstantBlue Coomassie Stain (Abcam).

### *In-vitro* kinase assays

All steady-state *in-vitro* kinase assays were assembled in 10µL volume in a buffer containing 25mM HEPES pH 7.5, 150mM NaCl, 1mM MgCl_2_, 2mM DTT, 5% glycerol, and 1mg/mL native BSA. All kinase assays also contained 25nM ATM kinase. The substrate used for all kinase assays is a GST-tagged p53 N-terminal 44aa fragment (GST-p53 residues 1-44). Reactions were assembled on ice, and started by the addition of 0.5mM cold ATP supplemented with 4μCi [γ-^32^P] ATP (6000 Ci/mmol, Perkin-Elmer) per reaction. Reactions proceeded for 1 hour at 30°C, and were stopped by the addition of 20µL of stop buffer containing 1.5X NuPAGE LDS sample buffer, 40mM EDTA, and 30mM TCEP. 10µL (1/3 of the reaction) was loaded onto a 4-12% Bis-Tris SDS-PAGE gel and proteins were separated by electrophoresis. ATP standards were spotted and gels were dried on DE-81 ion exchange cellulose paper. Dried gels were exposed to a phosphor imaging plate and imaged on a Typhoon FLA 7000 imager. Bands were quantified using ImageJ software and the concentrations phosphorylated substrate and enzymatic velocities were calculated based on spotted ATP standards. Velocities were normalized to the concentration of ATM (V/[E] (s^-1^)) and enzymatic parameters were calculated by fitting the data to the Michaelis-Menten equation:

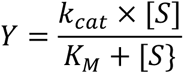

For MRN/dsDNA titrations and EC_50_ calculations (Figure 5A to C, Supplemental Figure 10C, Supplemental Figure 11B and C) the substrate was kept fixed at 25µM concentration. EC_50_ values, hill coefficients (*n*), and fold-activation were calculated by fitting the data to the following “[agonist] vs. response (four parameters)” equation in Prism software:

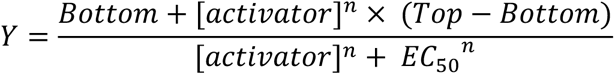

For single-turnover assays (Supplemental Figure 11G), large reactions were assembled containing 250nM ATM + 25nM 250bp dsDNA ± 250nM MRN ± 100nM p53 substrate and incubated at 25°C for 5 minutes. ATP was spiked in to a final concentration of 0.5mM containing 4μCi [γ-^32^P] ATP and reactions were incubated at 25°C. 10µL aliquots were removed at specified time points and mixed with 20µL of stop buffer. The remainder of the assay was performed in the same manner as steady-state kinase assays.

### Data availability

The refined dimer structures and corresponding cryo-EM maps (including consensus, symmetry expanded, focused, and composite maps) for unbound and Nc28-bound ATM have been deposited within the Protein Data Bank (PDB) and the Electron Microscopy Data Bank (EMDB) under accession codes 7SIC and EMD-25140 (unbound) and 7SID and EMD-25141 (Nc28-bound).

## Acknowledgements

We thank Haijuan Yang for establishing the FLAG-tagged ATM HEK 293 stable cell line and establishing preliminary purification procedures. We also thank Jason De La Cruz and Doreen Matthies for assistance in data collection at the MSK and HHMI cryoEM facilities, respectively. This work was supported by HHMI and NIH grant CA008748 (to N.P.) and NCI training grant 5F32CA247320 (to C.W.).

## Supplemental figure legends

**Supplemental Figure 1.**
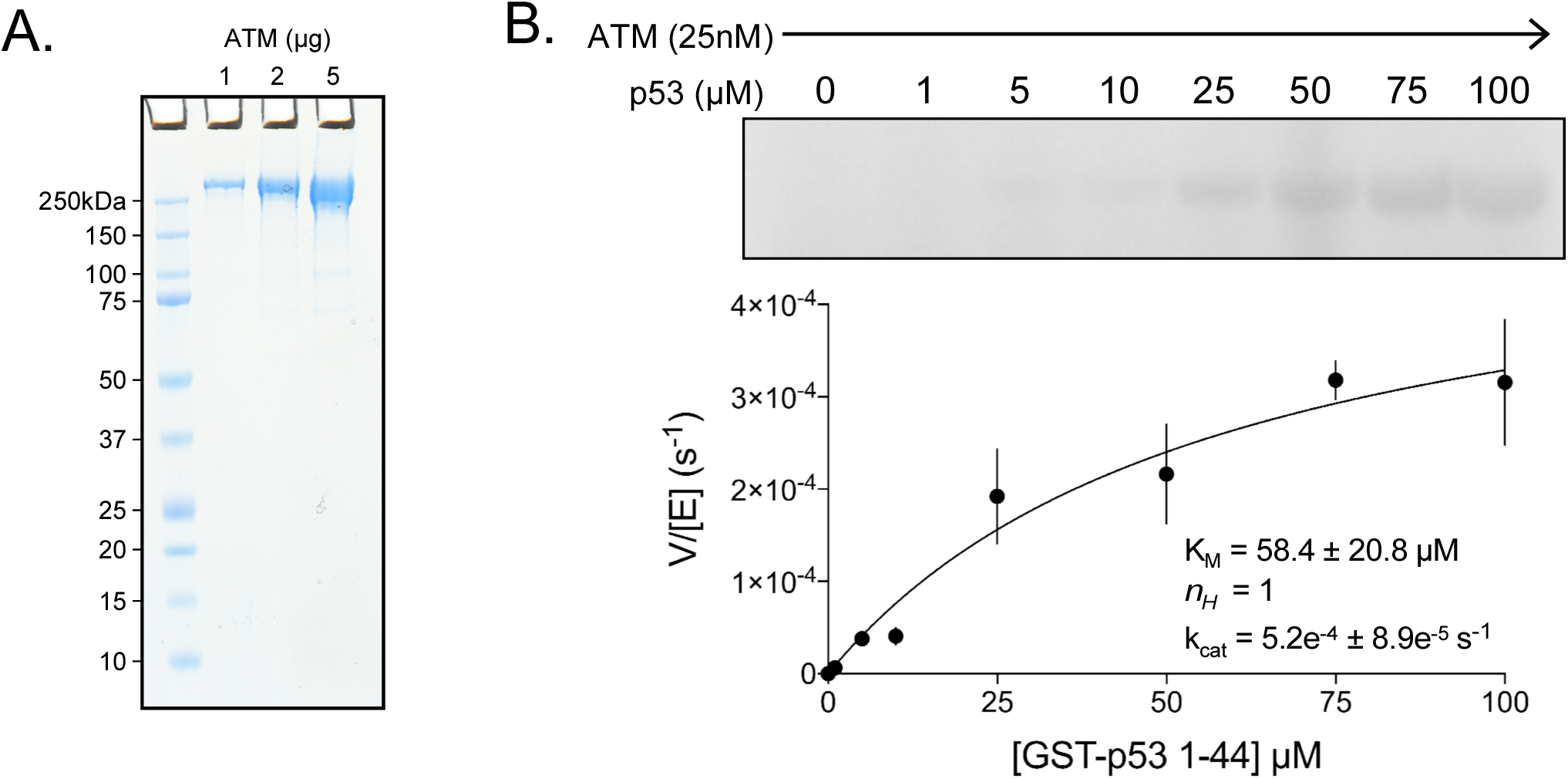
Purification and basal activity of ATM kinase. **A)** SDS-PAGE of purified FLAG-tagged ATM kinase. 1, 2 and 5 µg of protein were run on the gel. **B)** <u>Top</u>: Steady-state kinase assay of ATM with various concentrations of GST-tagged p53 N-terminal 44 residue substrate (GST-p53 1 to 44) with no dsDNA or MRN added. <u>Bottom</u>: Quantification of enzymatic parameters (K_M_, *n_H_*, and k_cat_) of ATM toward GST-tagged p53 peptide substrate. 3 biological replicates per point. Curve fit to the Michaelis-Menten equation.

**Supplemental Figure 2.**
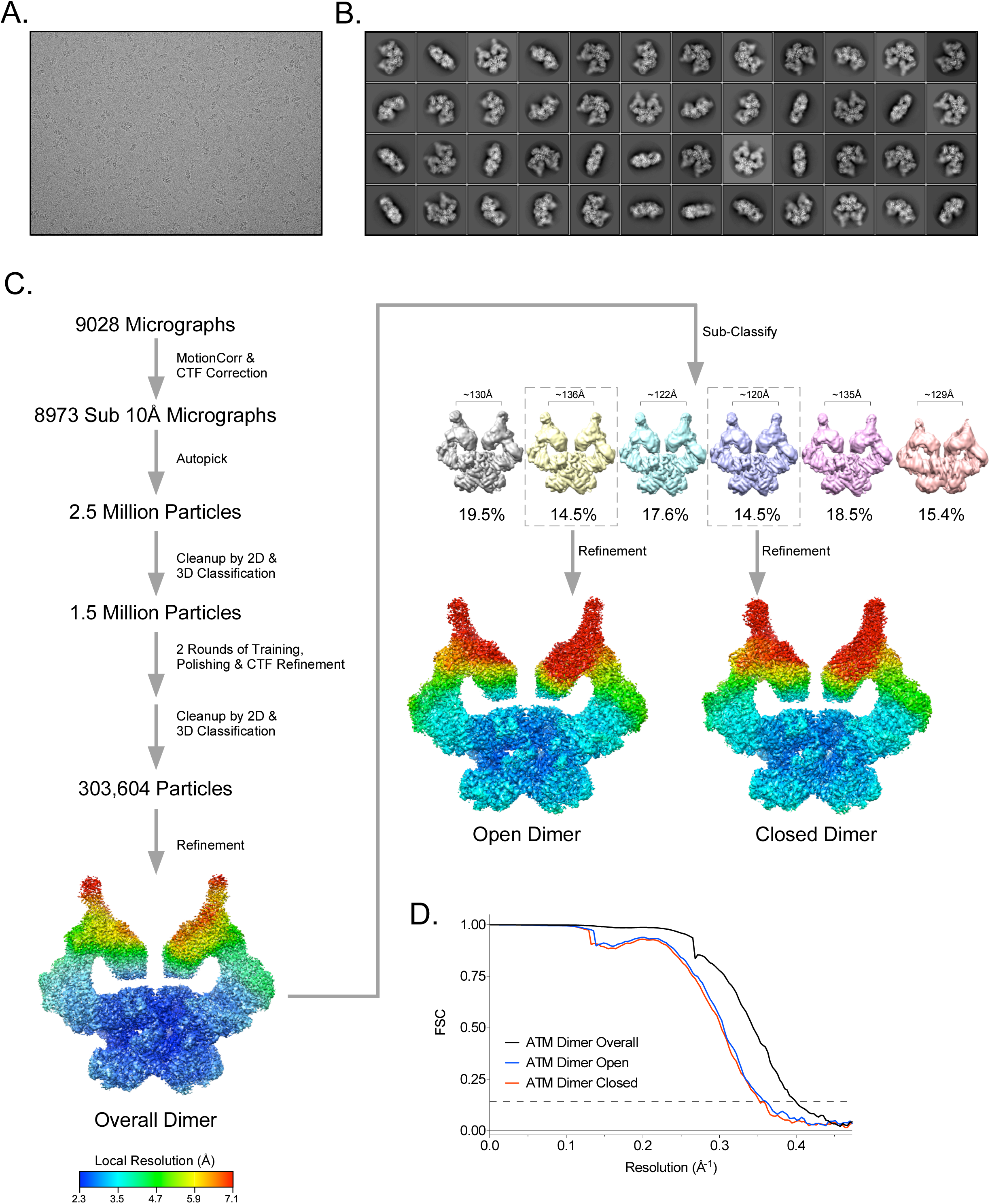
Cryo-EM data processing of the apo ATM sample. **A)** Example micrograph of ATM. **B)** Example 2D class averages of ATM. **C)** Workflow of ATM cryo-EM data analysis. Rainbow maps are colored by local resolution, color scale at bottom left. **D)** Fourier shell correlation (FSC) curves of B-factor sharpened overall, open, and closed ATM dimer maps. Gold-standard resolution threshold (FSC=0.143) shown as a dashed line.

**Supplemental Figure 3.**
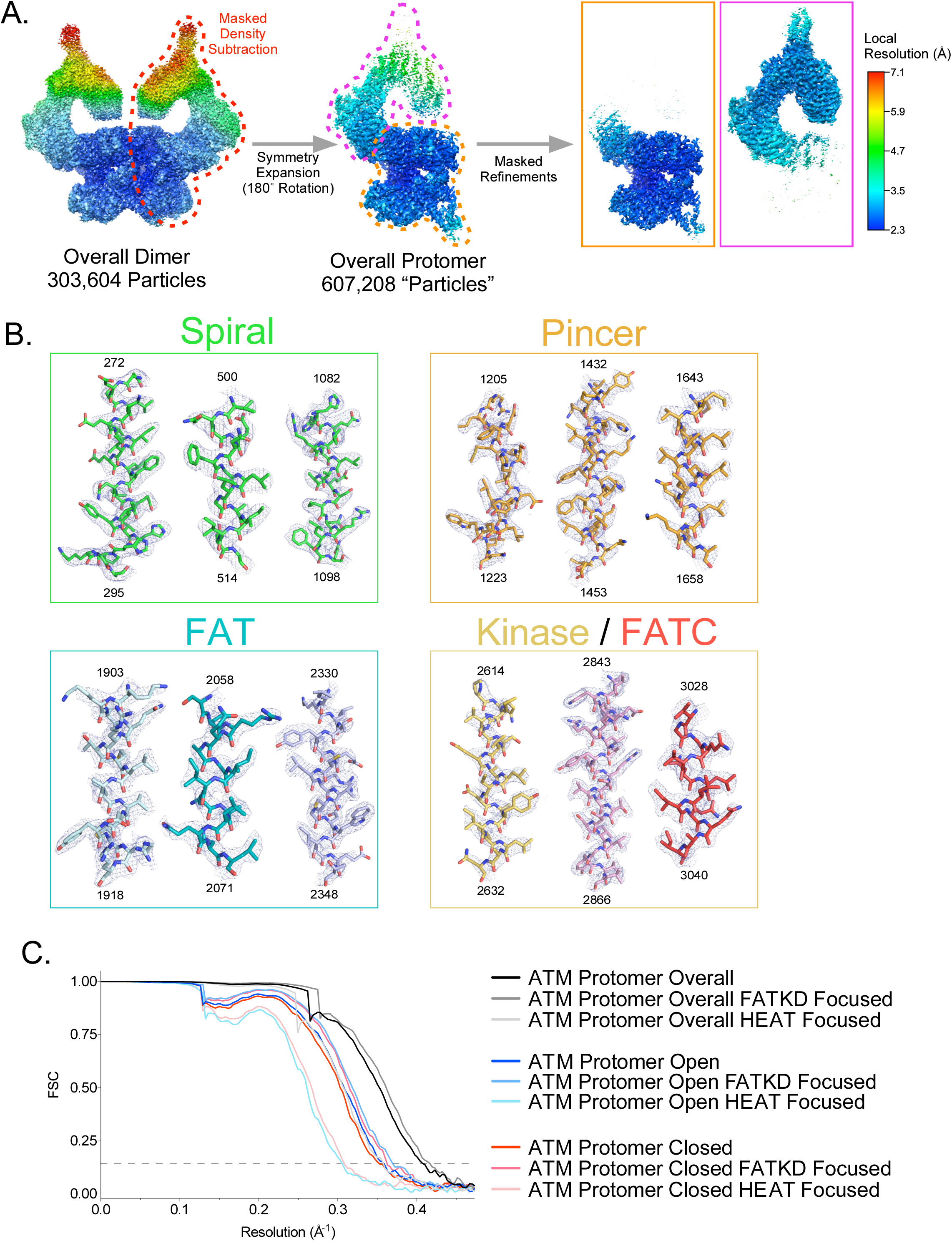
Partial signal subtraction, symmetry expansion and focused refinement procedure for ATM. **A)** Graphic of the partial signal subtraction, symmetry expansion and focused refinement procedure. FATKD and HEAT-focused refinement maps shown in orange and magenta boxes on right, respectively. All maps colored by local resolution based on the scale on the right. **B)** Electron density snapshots of helices within the Spiral, Pincer, FAT, and Kinase/FATC domains. Helices colored according to domain as in Figure 1A. Corresponding residue numbers are displayed at the top and bottom of each helix. All maps are contoured to 7σ. **C)** FSC curves of B-factor sharpened overall, open, and closed protomer maps, including FATKD and HEAT focused maps for each. Gold-standard resolution threshold (FSC=0.143) shown as a dashed line.

**Supplemental Figure 4.**
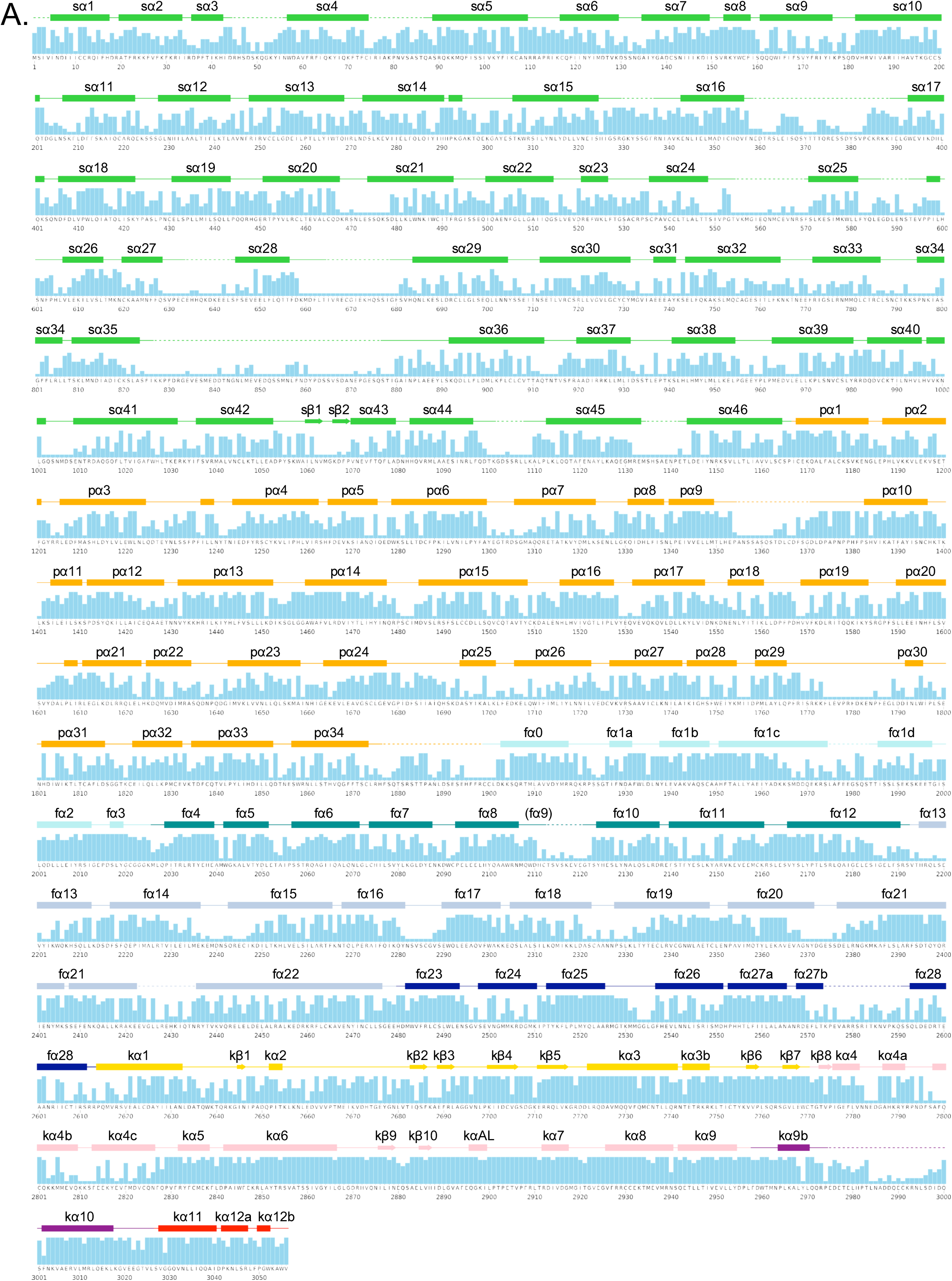
Secondary structure and sequence conservation of human ATM. **A)** Sequence of human ATM kinase showing CONSURF scores as bar graph for each residue. STRIDE-calculated secondary structures of the overall protomer are mapped onto the sequence. Helices are shown as rectangles, sheets as arrows, and coils as solid lines. Disordered regions not built in our structure are shown as dashed lines. Secondary structures are colored by domain as in Figure 1A and labeled.

**Supplemental Figure 5.**
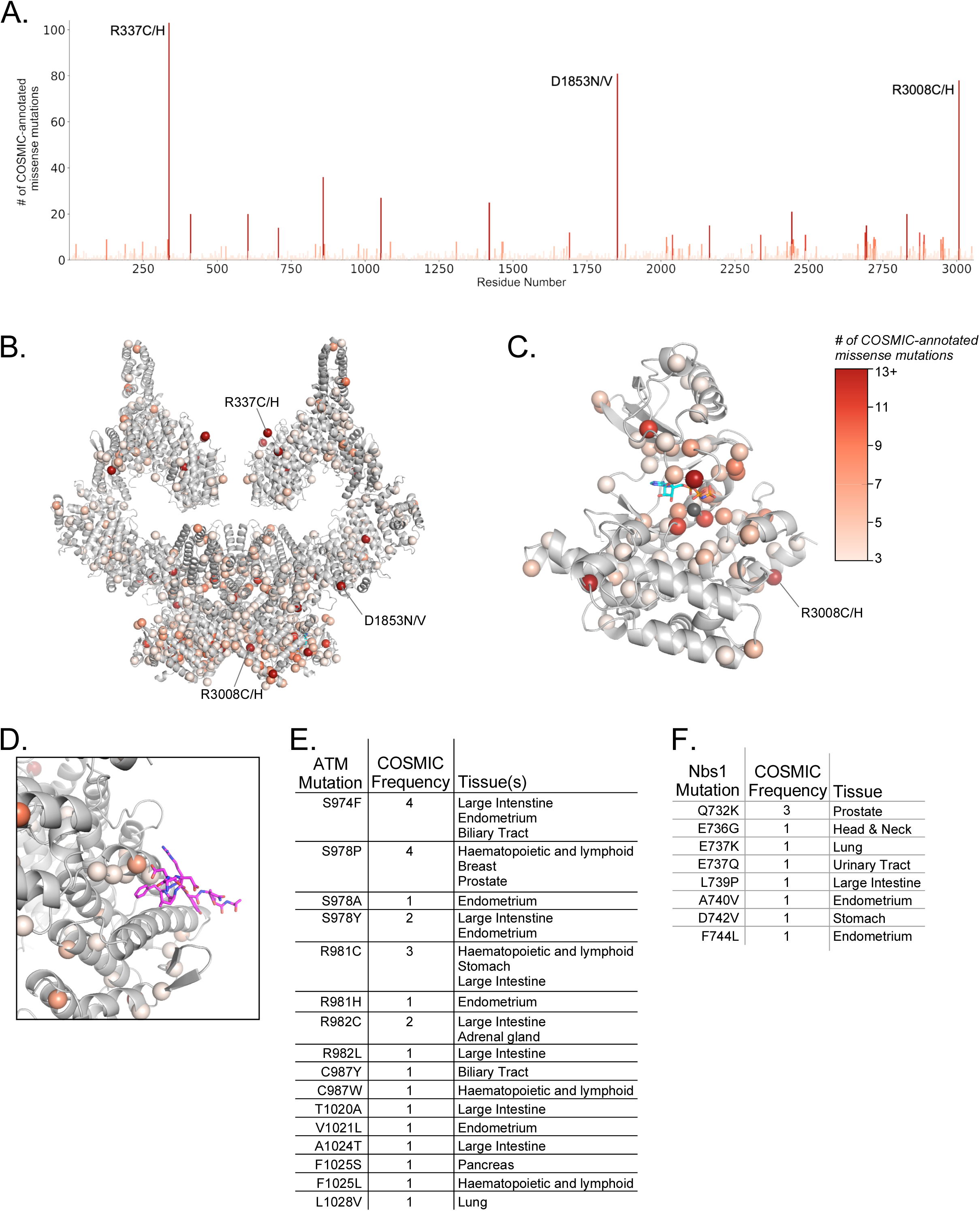
Positions and frequencies of cancer-associated ATM missense mutations. **A)** Positions and frequencies of cancer associated missense mutations from the COSMIC database along the sequence of ATM. The three most prevalent missense mutations are labeled. **B)** Positions of cancer associated missense mutations from the COSMIC database mapped onto the overall ATM dimer structure as spheres. Frequencies of each mutation shown by color scale (right of figure). Positions with less than 3 COSMIC-annotated mutations are not displayed for clarity. **C)** Positions and frequencies of cancer associated missense mutations within the ATM kinase domain. AMP-PNP and magnesium ion colored cyan and gray, respectively. **D)** Positions and frequencies of ATM cancer associated missense mutations within the Nbs1 binding cleft. Nbs1 C-terminal peptide (Nc28) shown as magenta sticks. **E)** Table of ATM missense mutations, frequencies, and associated cancer types that may interfere with Nbs1 binding. **F)** Table of all Nbs1 missense mutations, frequencies, and associated cancer types from the C-terminal 28 amino acids.

**Supplemental Figure 6.**
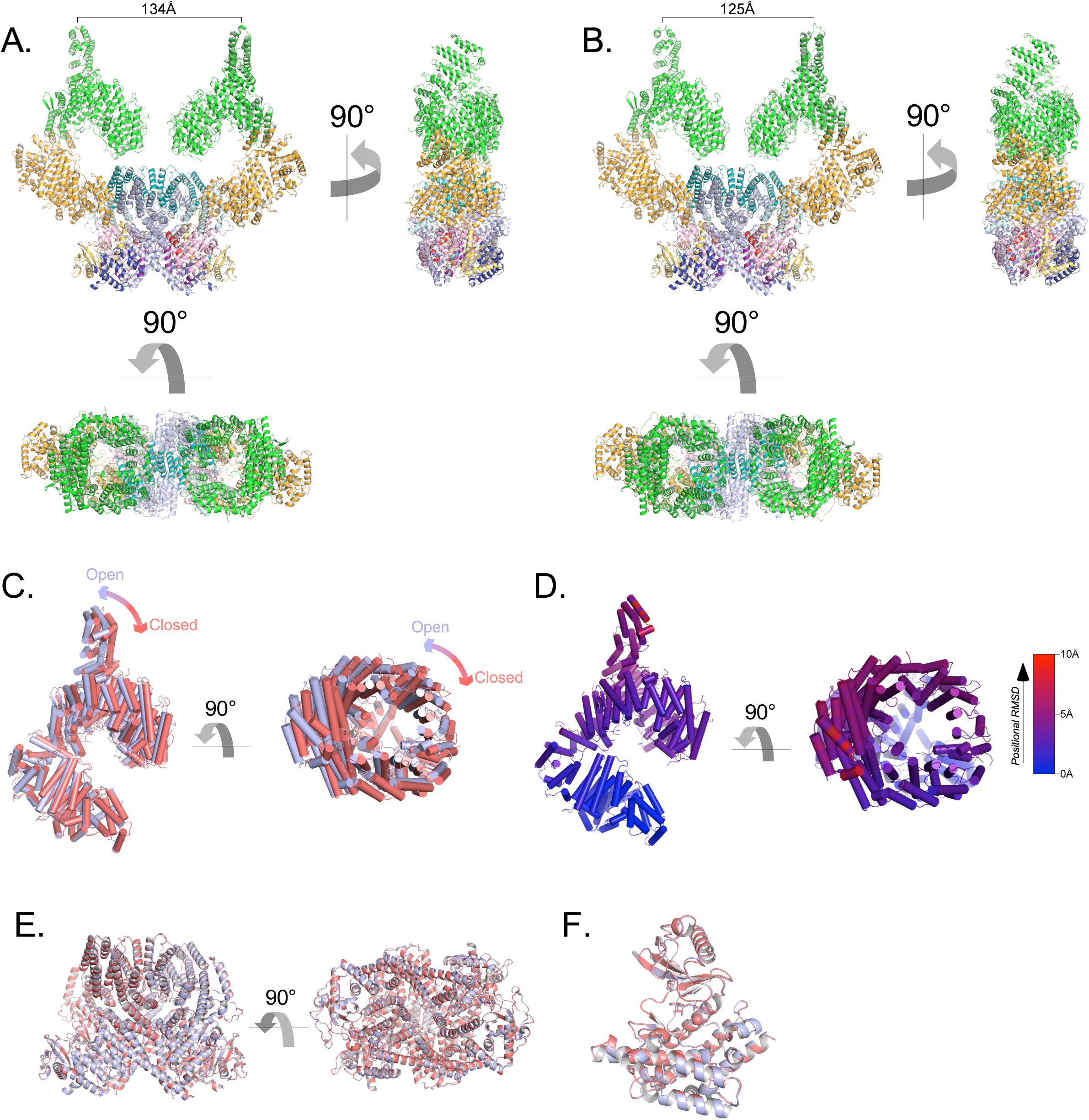
Flexibility of the ATM spiral and pincer domains. **A)** Structure of the most open class of the ATM dimer. Cα-Cα distance between protomers from residue Ile10 on sα1 shown. **B)** Structure of the most closed class of the ATM dimer. Cα-Cα distance between protomers from residue Ile10 on sα1 shown. **C)** Alignment of the Spiral and Pincer domains of the most open and closed ATM protomers. Open structure is colored light blue, closed structure is colored red. **D)** Structure of the Spiral and Pincer domains of the most open ATM protomer colored by Cα positional RMSD (0-10Å, blue to red) compared to the closed protomer. Generated by the ColorByRMSD PyMOL script. **E)** Alignment of the FATKD of the most open and most closed classes. Overall Cα positional RMSD = 0.39 Å. **F)** Alignment of the kinase domains of the most open and closed classes. Overall Cα positional RMSD = 0.24 Å.

**Supplemental Figure 7.**
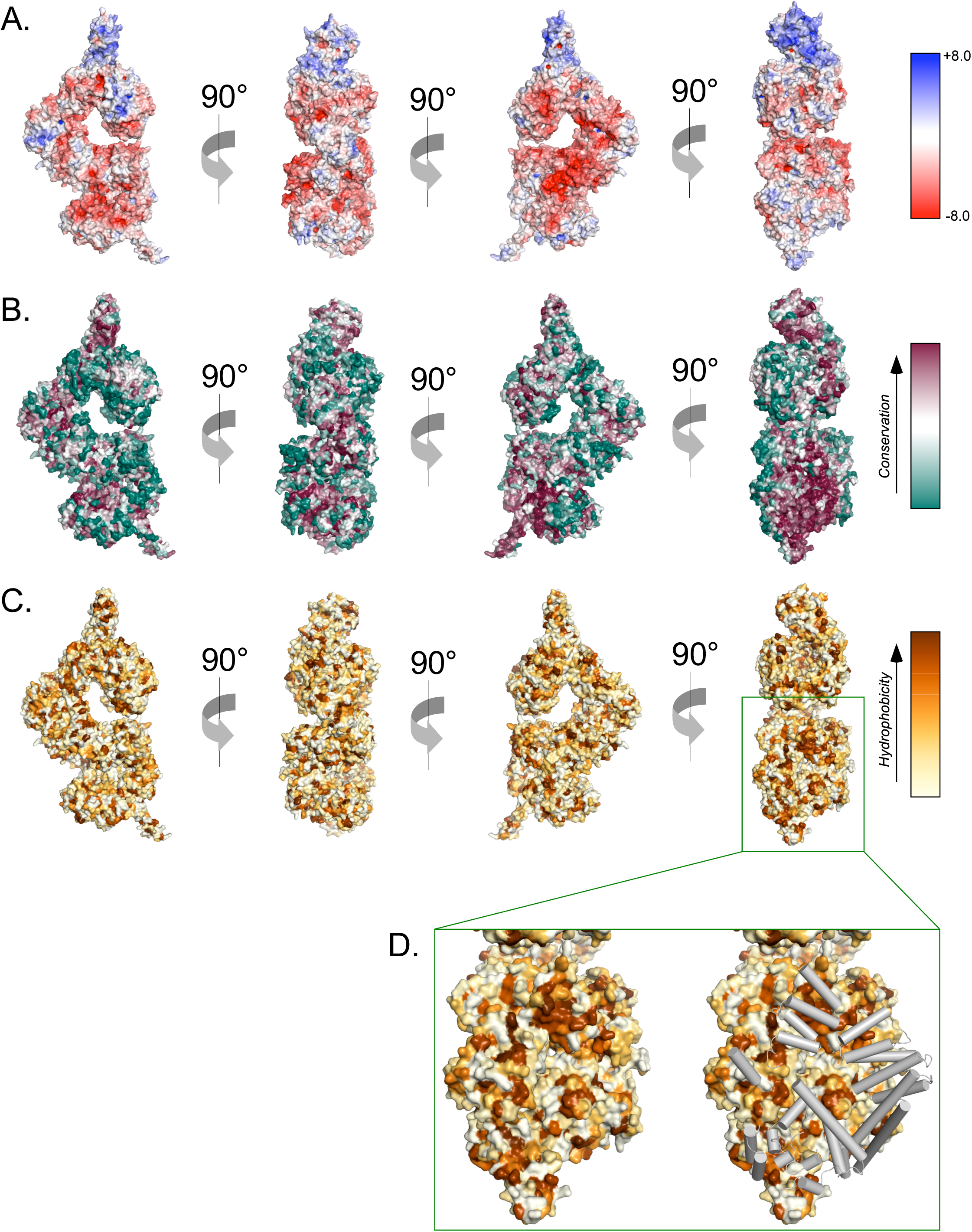
Surface properties of the ATM protomer. **A)** Overall ATM protomer colored by APBS-calculated surface potential. Charge scale shown on the right. **B)** Overall ATM protomer colored by CONSURF sequence conservation scores. Conservation scale shown on the right. **C)** Overall ATM protomer colored by relative hydrophobicity. Relative hydrophobicity scale shown on the right. **D)** <u>Left</u>: zoom on the dimer interface including FAT, Kinase, and FATC domains showing multiple large hydrophobic patches. <u>Right</u>: Identical view showing FAT, Kinase, and FATC domain helices from the symmetric protomer as gray cylinders that shield hydrophobic patches in the dimer structure.

**Supplemental Figure 8.**
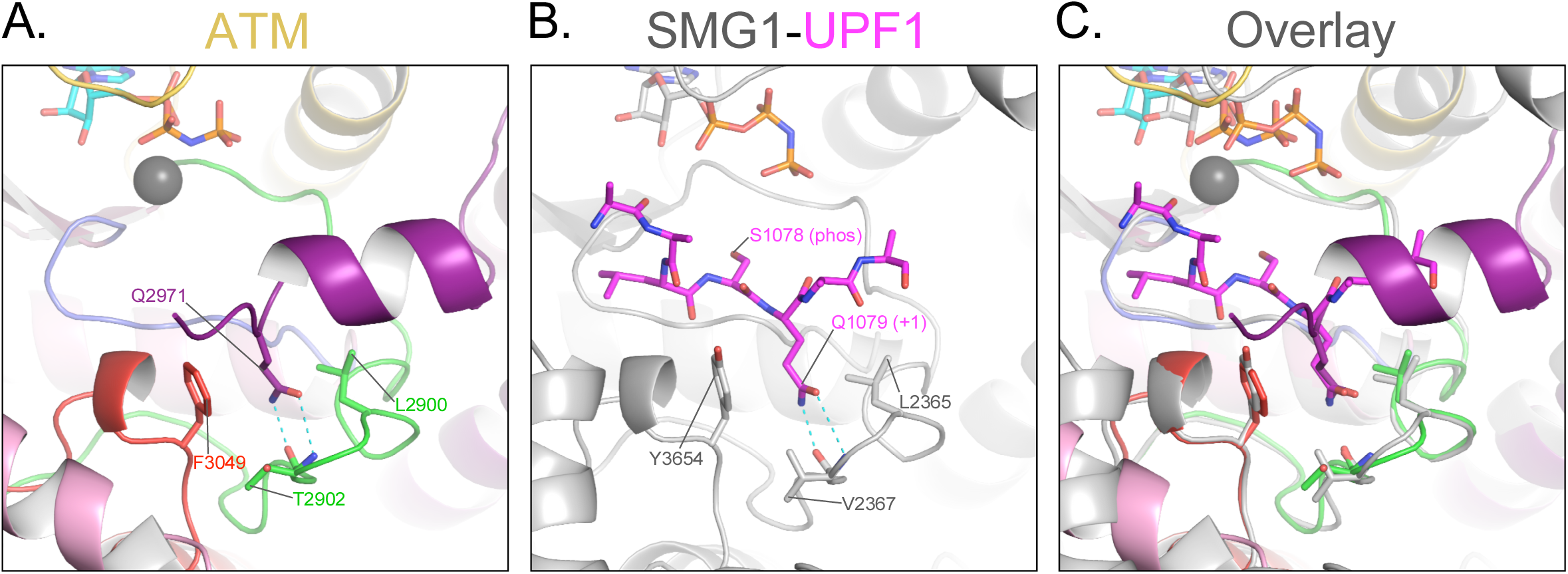
Blockage of the substrate-binding site. **A)** Structure of ATM catalytic cleft showing Gln2971 hydrogen bonding to backbone amide and carbonyl groups of Thr2902 on the activation loop, anchored between hydrophobic residues Leu2900 and Phe3049. **B)** Structure of Smg1 catalytic cleft bound to a Upf1 substrate peptide showing Gln1079 in the P_+1_ position hydrogen bonding to backbone amide and carbonyl groups of Val2367 on the activation loop, anchored between hydrophobic residues Leu2365 and Tyr3654. **C)** Alignment of the activation loops of ATM and Smg1 showing the blockage of the substrate binding site by ATM Gln2971.

**Supplemental Figure 9.**
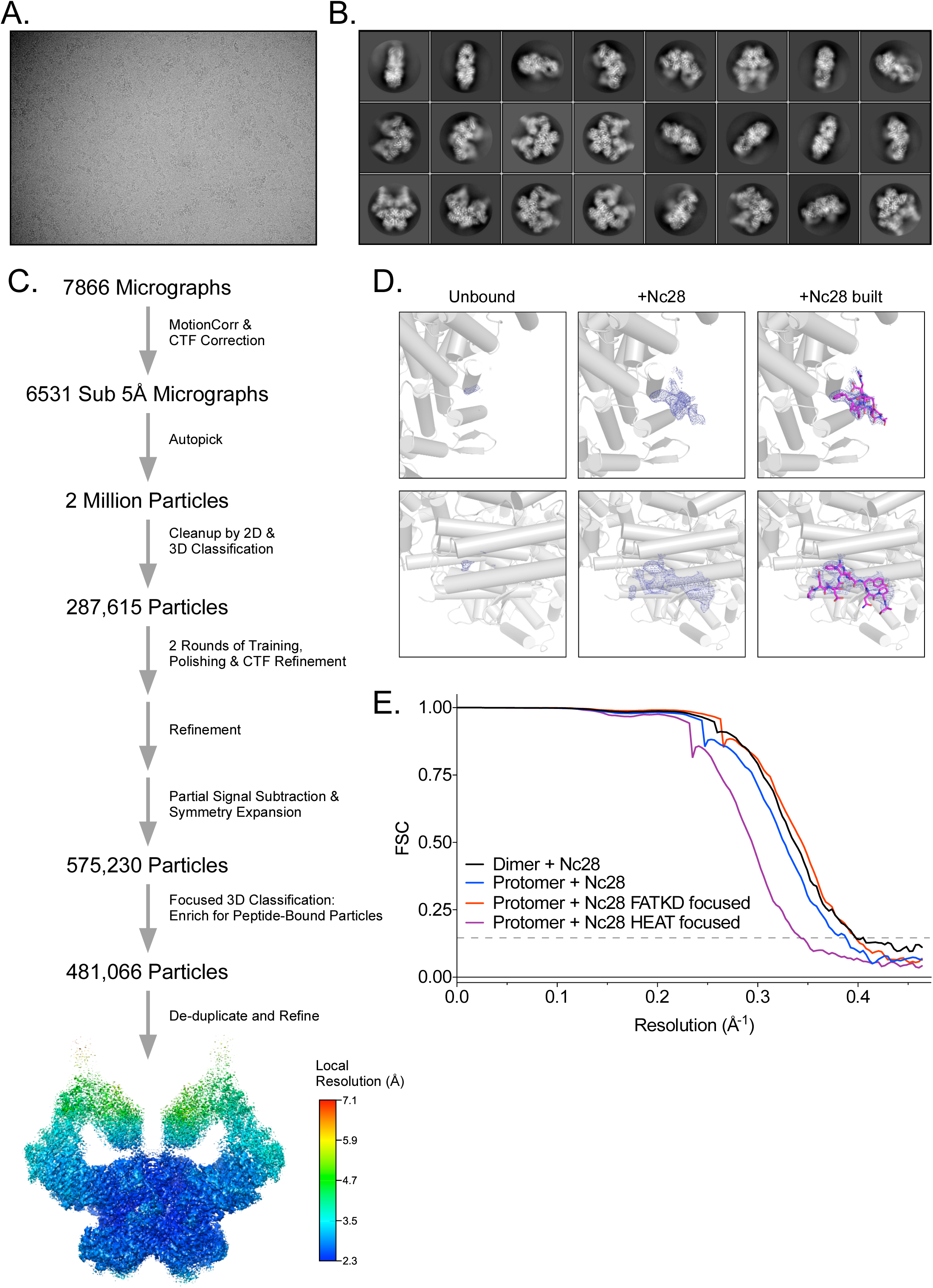
Cryo-EM data processing of the ATM-Nc28 sample. **A)** Example micrograph of ATM-Nc28. **B)** Example 2D class averages of ATM-Nc28. **C)** Workflow of ATM-Nc28 cryo-EM data analysis. Rainbow consensus map is colored by local resolution, color scale on the right. **D)** Electron density within the hydrophobic pocket of ATM without and with the Nc28 peptide in the sample. Rows show two different views of the hydrophobic pocket. First column of images shows minimal electron density in this region in the HEAT-focused overall apo map (607,208 particles). Second column shows much stronger density in this region in the HEAT-focused ATM-Nc28 map (481,066 particles). Third column shows the model of the Nc28 peptide structure built into the density shown in the second column. All maps are contoured to 6σ. **E)** Fourier shell correlation (FSC) curves of the ATM-Nc28 dimer and protomer maps, including FATKD and HEAT-focused maps. Gold-standard resolution threshold (FSC=0.143) shown as a dashed line.

**Supplemental Figure 10.**
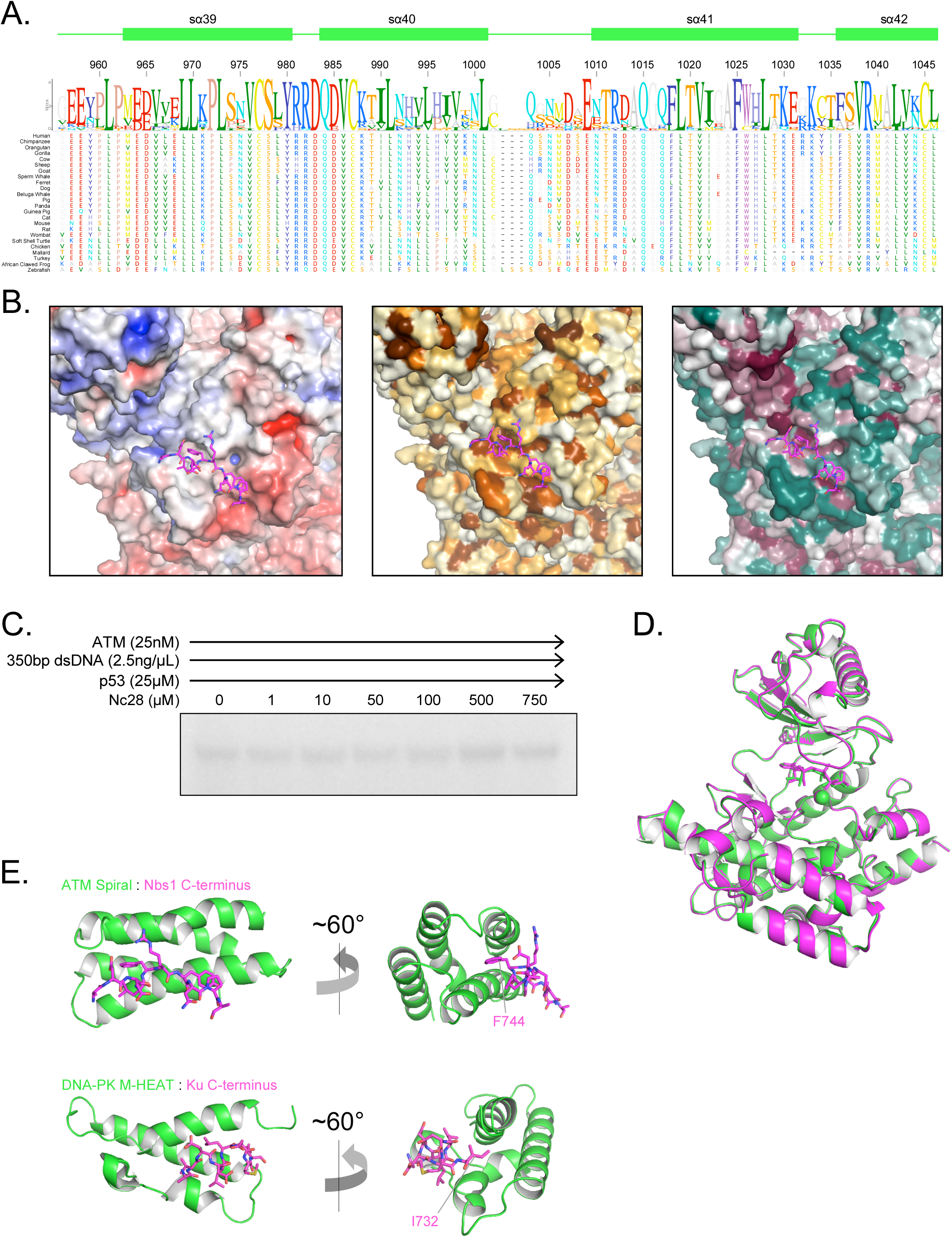
Structural assessment of ATM bound to the Nc28 peptide. **A)** Alignment and sequence conservation of ATM Spiral domain residues 956-1046 from 24 vertebrate species. Residue numbers and secondary structures are displayed according to the human ATM. **B)** Nbs1 peptide binding site shown as surface representation colored by APBS-calculated surface potential (left), hydrophobicity (middle), or CONSURF score (right). Same coloring scheme used as in Supplemental Figure 7. **C)** Steady-state ATM kinase assay using 25nM ATM, 2.5ng/µL 350bp dsDNA, 25µM p53 substrate, and various concentrations of the Nc28 peptide showing no increase in catalytic activity. Reactions were performed 3 times with similar results. **D)** Structural alignment of the kinase domains of apo (green) and Nc28-bound (magenta) ATM. Overall Cα positional RMSD = 0.29 Å. **E)** Comparison of the structures of Nc28 bound to the ATM Spiral domain (top) and the C-terminus of Ku bound to the DNA-PKcs M-HEAT domain (bottom).

**Supplemental Figure 11.**
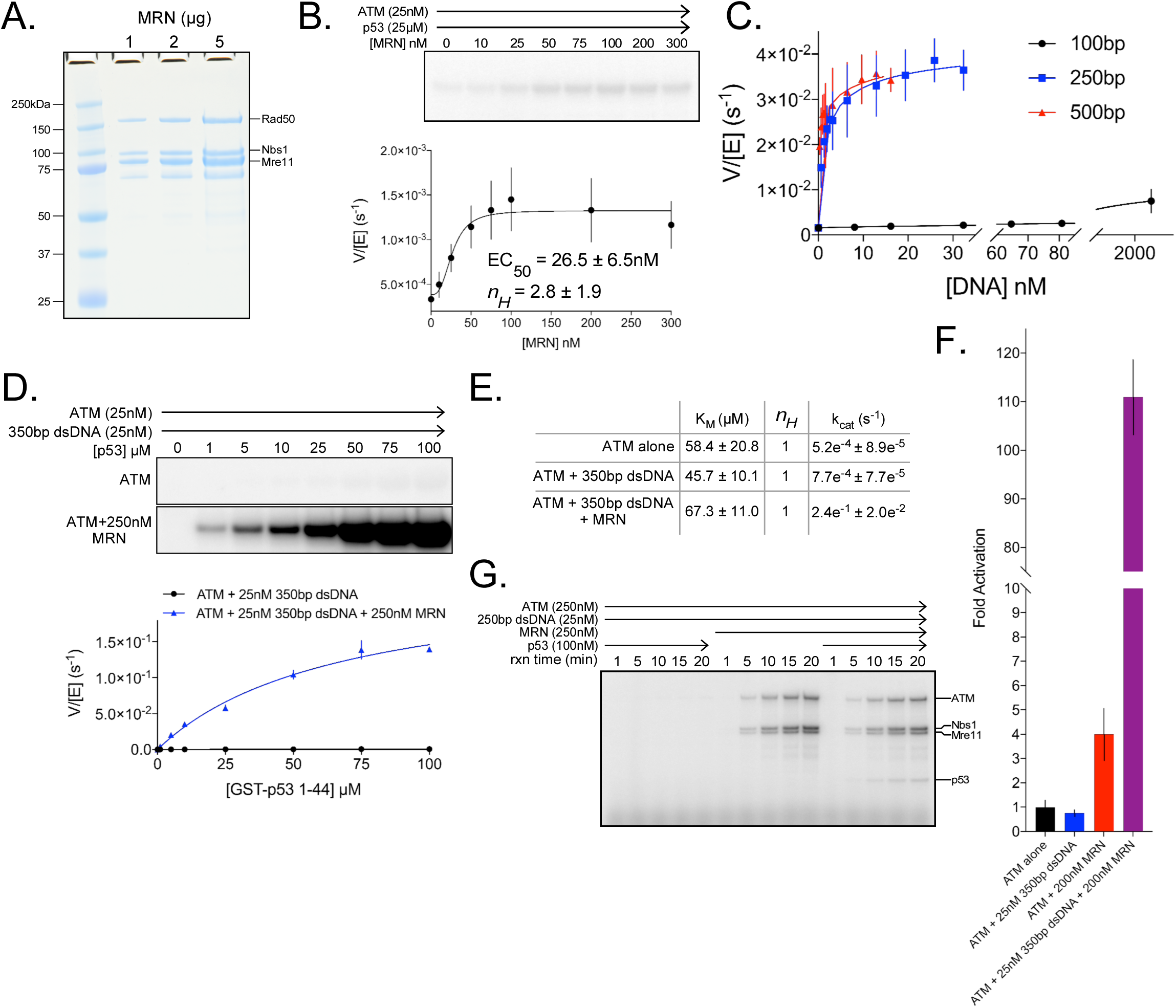
Quantitative ATM kinase assays. **A)** SDS-PAGE of purified FLAG-tagged MRN. 1, 2 and 5 µg of protein were run on the gel. **B)** <u>Top:</u> Steady-state kinase assay using 25 nM ATM, 25 µM p53 substrate, and various concentrations of MRN. <u>Bottom:</u> Quantification of ATM enzymatic velocity as a function of MRN concentration. 3 biological replicates per point. Curve fit to the EC_50_ equation. **C)** Quantification of ATM enzymatic velocity as a function of DNA concentration. Same data used as in Figure 5B with the x-axis rescaled to molar DNA concentration. 3 biological replicates per point, Curve fit to the EC_50_ equation. **D)** <u>Top:</u> Steady-state kinase assay using 25 nM ATM, 25 nM 350bp dsDNA, and various concentrations of p53 substrate peptide without and with 250 nM MRN added. <u>Bottom:</u> Quantification of ATM enzymatic velocity as a function of substrate concentration without (black) and with (blue) MRN present. 3 biological replicates per point, Curve fit to the Michaelis-Menten equation. **E)** Quantification of enzymatic parameters (K_M_ and k_cat_) of ATM under various conditions from data in Supplemental Figures 1B and 11D. **F)** Data from Supplemental Figures 1B, 11B, and 11D rescaled to show fold activation relative to ATM alone at 25 µM substrate concentration. 3 biological replicates per group. **G)** Time-course kinase assay run under single-turnover conditions using 250 nM ATM + 25 nM 250bp dsDNA ± 250 nM MRN ± 100 nM p53 substrate peptide. Phosphorylation products are labeled on the right. Reactions were performed 3 times with similar results.

**Supplemental Figure 12.**
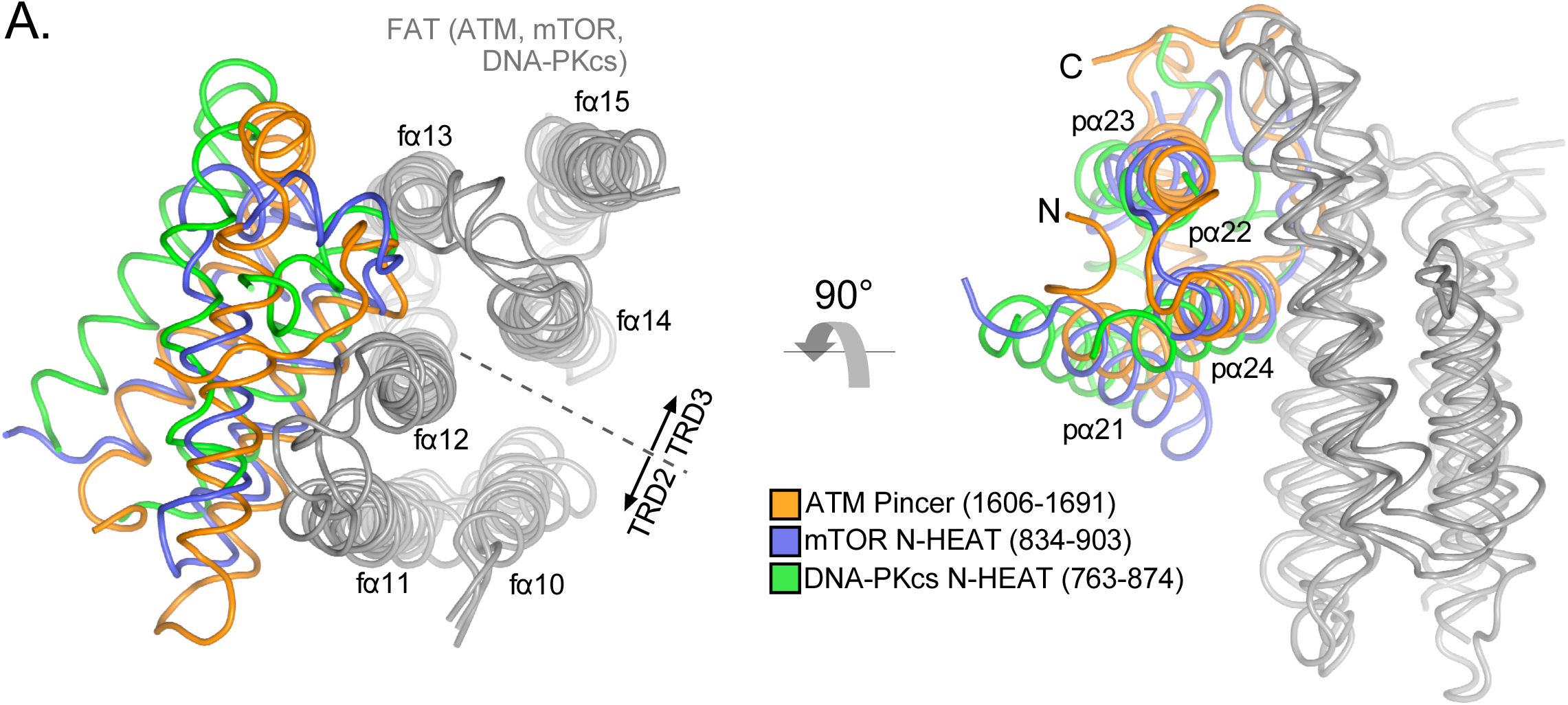
Structural conservation of the FAT domain anchor. **A)** Alignment of the TRD2-TRD3 interface helices from ATM, mTOR, and DNA-PKcs inactive structures (gray). ATM FAT domain helices fα10 to fα15 are labeled on the left. ATM Pincer helices pα21 to pα24 labeled on the right. Four helix bundles from the mTOR and DNA-PKcs N-HEAT domains are shown in blue and green, respectively.

**Supplemental Table 1.**
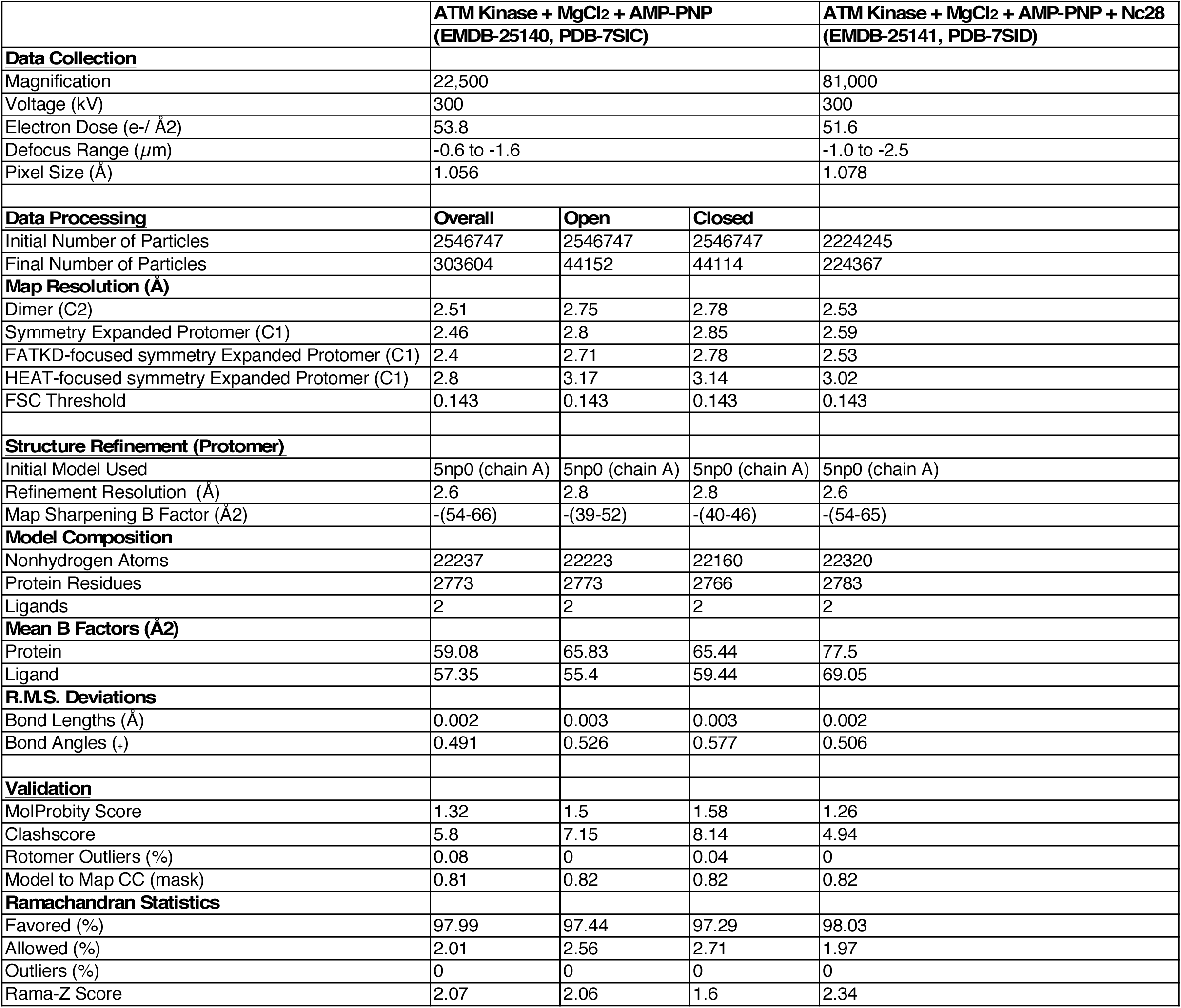
Cryo-EM data processing and refinement table.

|  | ATM Kinase + MgCl2 + AMP-PNP<br>(EMDB-25140, PDB-7SIC) |  |  | ATM Kinase + MgCl2 + AMP-PNP + Nc28<br>(EMDB-25141, PDB-7SID) |
| --- | --- | --- | --- | --- |
| Data Collection |  |  |  |  |
| Magnification | 22,500 |  |  | 81,000 |
| Voltage (kV) | 300 |  |  | 300 |
| Electron Dose (e-/ Å2) | 53.8 |  |  | 51.6 |
| Defocus Range (μm) | -0.6 to -1.6 |  |  | -1.0 to -2.5 |
| Pixel Size (Å) | 1.056 |  |  | 1.078 |
| Data Processing | Overall | Open | Closed |  |
| Initial Number of Particles | 2546747 | 2546747 | 2546747 | 2224245 |
| Final Number of Particles | 303604 | 44152 | 44114 | 224367 |
| Map Resolution (Å) |  |  |  |  |
| Dimer (C2) | 2.51 | 2.75 | 2.78 | 2.53 |
| Symmetry Expanded Protomer (C1) | 2.46 | 2.8 | 2.85 | 2.59 |
| FATKD-focused symmetry Expanded Protomer (C1) | 2.4 | 2.71 | 2.78 | 2.53 |
| HEAT-focused symmetry Expanded Protomer (C1) | 2.8 | 3.17 | 3.14 | 3.02 |
| FSC Threshold | 0.143 | 0.143 | 0.143 | 0.143 |
| Structure Refinement (Protomer) |  |  |  |  |
| Initial Model Used | 5np0 (chain A) | 5np0 (chain A) | 5np0 (chain A) | 5np0 (chain A) |
| Refinement Resolution (Å) | 2.6 | 2.8 | 2.8 | 2.6 |
| Map Sharpening B Factor (Å2) | -(54-66) | -(39-52) | -(40-46) | -(54-65) |
| Model Composition |  |  |  |  |
| Nonhydrogen Atoms | 22237 | 22223 | 22160 | 22320 |
| Protein Residues | 2773 | 2773 | 2766 | 2783 |
| Ligands | 2 | 2 | 2 | 2 |
| Mean B Factors (Å2) |  |  |  |  |
| Protein | 59.08 | 65.83 | 65.44 | 77.5 |
| Ligand | 57.35 | 55.4 | 59.44 | 69.05 |
| R.M.S. Deviations |  |  |  |  |
| Bond Lengths (Å) | 0.002 | 0.003 | 0.003 | 0.002 |
| Bond Angles (.) | 0.491 | 0.526 | 0.577 | 0.506 |
| Validation |  |  |  |  |
| MolProbity Score | 1.32 | 1.5 | 1.58 | 1.26 |
| Clashscore | 5.8 | 7.15 | 8.14 | 4.94 |
| Rotomer Outliers (%) | 0.08 | 0 | 0.04 | 0 |
| Model to Map CC (mask) | 0.81 | 0.82 | 0.82 | 0.82 |
| Ramachandran Statistics |  |  |  |  |
| Favored (%) | 97.99 | 97.44 | 97.29 | 98.03 |
| Allowed (%) | 2.01 | 2.56 | 2.71 | 1.97 |
| Outliers (%) | 0 | 0 | 0 | 0 |
| Rama-Z Score | 2.07 | 2.06 | 1.6 | 2.34 |

